# Site of vulnerability on SARS-CoV-2 spike induces broadly protective antibody to antigenically distinct omicron SARS-CoV-2 subvariants

**DOI:** 10.1101/2022.10.31.514592

**Authors:** Siriruk Changrob, Peter J. Halfmann, Hejun Liu, Jonathan L. Torres, Joshua J.C. McGrath, Gabriel Ozorowski, Lei Li, Makoto Kuroda, Tadashi Maemura, Min Huang, G. Dewey Wilbanks, Nai-Ying Zheng, Hannah L. Turner, Steven A. Erickson, Yanbin Fu, Gagandeep Singh, Florian Krammer, D. Noah Sather, Andrew B. Ward, Ian A. Wilson, Yoshihiro Kawaoka, Patrick C. Wilson

**Affiliations:** Drukier Institute for Children’s Health, Weill Cornell Medicine, New York, New York, USA; Influenza Research Institute, Department of Pathobiological Sciences, School of Veterinary Medicine, University of Wisconsin-Madison, Madison, WI 53711; Department of Integrative Structural and Computational Biology, The Scripps Research Institute, La Jolla, CA 92037, USA; University of Chicago Department of Medicine, Section of Rheumatology, Chicago, IL 60637, USA; Division of Virology, Department of Microbiology and Immunology, Institute of Medical Science, University of Tokyo, 108-8639 Tokyo, Japan; Department of Pathology, Molecular and Cell Based Medicine, Icahn School of Medicine at Mount Sinai, New York, NY, USA; Center for Vaccine Research and Pandemic Preparedness, Icahn School of Medicine at Mount Sinai, New York, NY, USA; Center for Global Infectious Disease Research, Seattle Children’s Research Institute, Seattle, Washington, USA; Department of Pediatrics, University of Washington, Seattle, Washington, USA; Department of Global Health, University of Washington, Seattle, Washington, USA; The Skaggs Institute for Chemical Biology; The Scripps Research Institute; La Jolla, CA 92037; USA

## Abstract

The rapid evolution of SARS-CoV-2 Omicron variants has emphasized the need to identify antibodies with broad neutralizing capabilities to inform future monoclonal therapies and vaccination strategies. Herein, we identify S728-1157, a broadly neutralizing antibody (bnAb) targeting the receptor-binding site (RBS) and derived from an individual previously infected with SARS-CoV-2 prior to the spread of variants of concern (VOCs). S728-1157 demonstrates broad cross-neutralization of all dominant variants including D614G, Beta, Delta, Kappa, Mu, and Omicron (BA.1/BA.2/BA.2.75/BA.4/BA.5/BL.1). Furthermore, it protected hamsters against *in vivo* challenges with wildtype, Delta, and BA.1 viruses. Structural analysis reveals that this antibody targets a class 1 epitope via multiple hydrophobic and polar interactions with its CDR-H3, in addition to common class 1 motifs in CDR-H1/CDR-H2. Importantly, this epitope is more readily accessible in the open and prefusion state, or in the hexaproline (6P)-stabilized spike constructs, as compared to diproline (2P) constructs. Overall, S728-1157 demonstrates broad therapeutic potential, and may inform target-driven vaccine design against future SARS-CoV-2 variants.

## Introduction

Since the start of the pandemic in December 2019, the severe acute respiratory syndrome coronavirus 2 (SARS-CoV-2) virus has led to over 576 million cases of coronavirus disease 2019 (COVID-19) and over six million deaths globally. Although the rapid development and distribution of vaccines and therapeutics has curbed the impact of COVID-19 to an extent, the emergence of circulating variants of concern (VOCs) continues to represent a major threat due to the potential for further immune evasion and enhanced pathogenicity. The D614G variant was the earliest variant to emerge and became universally prevalent thereafter. In comparison to wildtype (WT), the D614G variant exhibited increased transmissibility rather than increased pathogenicity and was therefore unlikely to reduce efficacy of vaccines in clinical trials^1^. Between the emergence D614G and October 2021, four additional significant VOC evolved worldwide, including Alpha, Beta, Gamma, and Delta. Among these variants, Delta became a serious global threat as a result of its transmissibility, increased disease severity, and partial immune evasion as shown by the reduced ability of polyclonal serum and monoclonal antibodies (mAbs) to neutralize this strain^2-6^. Shortly afterwards, in November 2021, the Omicron variant was identified and announced as a novel VOC. This variant possessed the largest number of mutations to date and appeared to spread more rapidly than previous strains^7,8^. Currently, there are five major subvariant lineages of Omicron (BA.1, BA.2, BA.3, BA.4 and BA.5) leading to new COVID-19 cases, with BA.5 becoming dominant over BA.2 and accounting for most new cases in the United States at the time of writing. The Omicron variants can escape recognition by COVID-19 vaccine-associated immunity to varying extents, thereby significantly reducing the neutralizing potency of serum antibodies from convalescent and fully mRNA-vaccinated individuals^9^. Similarly, Omicron variants were able to escape binding of several Emergency Use-Authorization (EUA) therapeutic mAbs even though these had been previously shown to be effective against earlier VOCs^10,11^. Due to the lowered neutralization against Omicron and the continued threat of future VOCs, there is an urgent need to identify broad and potent neutralizing antibodies that can protect against diverse evolving SARS-CoV-2 lineages.

In this study, we identify a potent RBD-reactive monoclonal antibody from the peripheral blood of SARS-CoV-2 convalescent individual that effectively neutralize Alpha, Beta, Kappa, Delta, Mu, and Omicron variants (BA.1, BA.2, BA.2.75, BA.4, BA.5 and BL.1). This mAb, S728-1157, can reduce BA.1 Omicron viral titer *in vivo* and significantly reduced viral loads during wildtype and Delta infection. In terms of specificity, S728-1157 bound the receptor binding site (RBS) that is fully exposed when the RBD on the spike is in the up conformation. S728-1157 binds using motifs found in the CDR-H1 and CDR-H2 domains that are common to IGHV3-53/3-66 class 1 antibodies but also via extensive unique contacts with CDR-H3 to circumvent mutations in the variant virus spikes. This suggests that the rational design of future vaccine boosts covering Omicron variants should be modified to present stabilized spike in the up configuration to optimally induce class 1 mAbs that have similar CDR-H3 features.

## Results

### Isolation of RBD-reactive mAbs that exhibit diverse patterns of neutralization and potency

Before the spread of the Omicron variant, we previously characterized 43 mAbs targeting distinct epitopes on the spike protein, including the N-terminal domain (NTD), RBD, and subunit 2 (S2), although none were able to neutralize all existing SARS-CoV-2 variants at that time^12^. In the current study, an additional panel of RBD-reactive mAbs were expressed from three high-responder subjects who mounted robust anti-spike IgG responses, as defined previously (Table S1 and Table S2)^13^. Although the proportion of spike RBD-binding B cells was similar in high-responders as compared to mid- and low-responders (Figure 1a-c), heavy chain somatic hypermutation rates were significantly greater in the high-responder group (Figure 1d), suggesting that these subjects may have the highest potential to generate potent cross-reactive mAbs^13^. These antibodies were assayed for binding to key RBD mutants to identify their epitope classifications (Table S3)^14^. Among 14 RBD-reactive mAbs, we identified four class 2 mAbs, two class 3 mAbs, and eight unclassified mAbs that showed little to no reduction of binding against any key RBD mutants tested (Figure 1f). Class 2 and 3 RBD mAbs did not recognize a multivariant RBD mutant containing K417N/E484K/L452R/N501Y substitutions, an artificially designed RBD to include key mutations for virus escape^14,15^, nor cross-reactivity to the RBD of SARS-CoV-1 and Middle Eastern respiratory syndrome (MERS)-CoV (Figure 1f). Functionally, class 2 and 3 RBD mAbs potently neutralized D614G and Delta but neutralizing activity was limited against Beta, Kappa and Mu (Figure 1g). No class 2 or 3 antibodies assayed could neutralize any tested Omicron variant.

**Figure 1.**
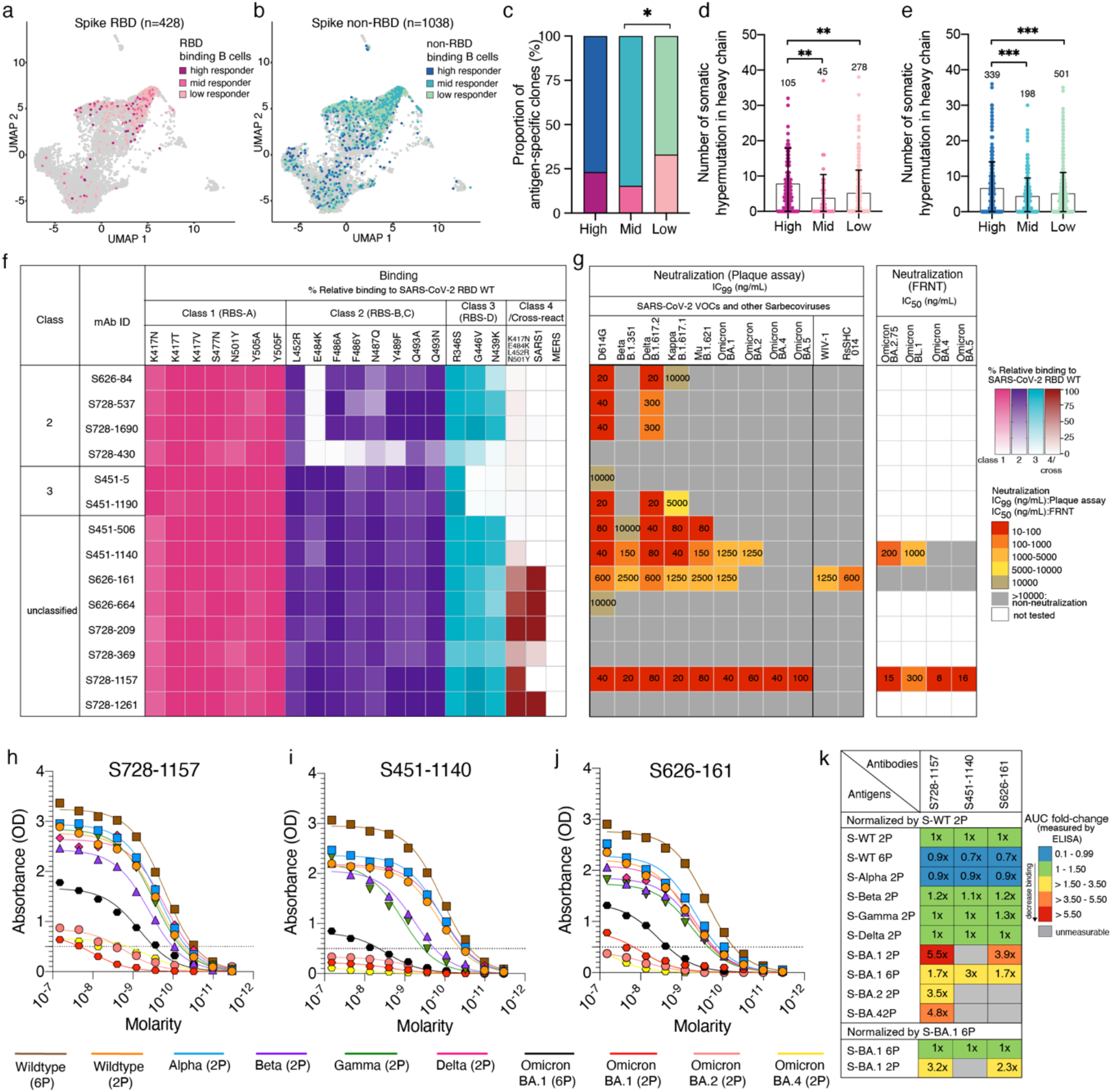
Proportion of SARS-CoV-2-specific B cells and characterization of RBD-reactive mAbs isolated from COVID-19 convalescent individuals. **a-b**, Uniform manifold approximation and projection (UMAP) of SARS-CoV-2 **(a)** spike RBD binding and **(b)** spike non-RBD binding B cells isolated from convalescent subjects that could be characterized into 3 groups (high, mid and low responder) based on their serological response against SARS-CoV-2 spike^13^. **c**, Proportion of spike non-RBD- and spike RBD-specific binding B cells representing in each responder group. Colors in **a** and **b** are representative of antigen-specific B cells from each responder group as follow: **(a)** RBD-binding B cells; plum, high responder; pink, mid responder; pale-pink, low responder and **(b)** non-RBD-binding B cells; navy, high responder; blue, mid responder; pale-green, low responder. **d-e**, Number of somatic hypermutations in the IGHV in antibodies targeting **(d)** RBD and **(e)** non-RBD. **f**, Binding profile of RBD-reactive mAbs against single RBD mutants associated with different antibody classes, a combinatorial RBD mutant, and the RBDs of SARS-CoV-1 and MERS-CoV. Color gradients indicate relative binding percentage compared to RBD WT with the labeling color as follow: pink, class 1; purple, class 2; teal, class 3; burgundy, class 4 and cross-reactive epitopes. **g**, Neutralization potency measured by plaque assay (complete inhibitory concentration; IC_99_) and focus reduction neutralization test (FRNT; half inhibitory concentration; IC_50_) of RBD-reactive mAbs to SARS-CoV-2 variants and sarbecoviruses. Binding breadth against full-length spike SARS-CoV-2 variants determined by ELISA is shown for **(h)** S728-1157, **(i)** S451-1140, and **(j)** S626-161. Dashed line in **h-j** indicate the limit of detection. **k**, Heatmap represents area under curve (AUC) fold-change of broadly neutralizing RBD-reactive mAbs against ectodomain spike SARS-CoV-2 variants relative to WT-2P and the differences of AUC fold-change between spike BA.1-2P relative to spike BA.1-6P. Colors in **k** indicate range of fold-change from blue, 0.1 to 0.99-fold (increase affinity binding); green, 1 to 1.5-fold (none to little reduction in affinity binding); yellow, >1.5 to 3.5-fold (moderate reduction in affinity binding); orange, >3.5 to 5.5-fold (high reduction in affinity binding); red, >5.5-fold (extreme reduction in affinity binding) and grey indicated unmeasurable fold-change due to the absorbance values are below limit of detection. The statistical analysis in **c** was determined using Tukey multiple pairwise-comparisons and in **d-e** was determined using Kruskal-Wallis with Dunn’s multiple comparison test. Data in **f-g and h-j** are representative of two independent experiments performed in duplicate. Genetic information for each antibody is in **Table S2**. The SARS-CoV-2 viruses used in neutralization assay are indicated in **Table S4**.

In contrast, the majority of unclassified mAbs bound to the RBD multivariant and cross-reacted to the SARS-CoV-1 RBD (Figure 1f). Among these, we went on to identify three bnAbs, S451-1140, S626-161 and S728-1157, that showed high neutralization potency against D614G and could cross-neutralize Beta, Delta, Kappa, Mu and BA.1 with 99% inhibitory concentration (IC_99_) in the range of 20-2500 ng/ml (Figure 1g). Given the broad neutralization potency of these three mAbs, in addition of plaque assay platform, we also performed the neutralization activity against authentic BA.2.75, BL.1 (BA.2.75+R346T), BA.4, and BA.5 viruses using focus reduction neutralization test (FRNT) (Figure 1g). Of these, S728-1157 displayed high neutralizing activities against the panel of Omicron variants including BA.1, BA.2, BA.4 and BA.5, with IC_99_ up to 100 ng/ml as measured by plaque assay. A similar scenario was observed using FRNT, S728-1157 maintains its high neutralization activity against BA.2.75, BL.1, BA.4 and BA.5 with 50% inhibitory concentration (IC_50_) in the range of 8-16 ng/ml (Figure 1g). S451-1140 neutralized BA.1, BA.2, BA.2.75 and BL.1 potently, but not BA.4 and BA.5 as observed in both neutralization assay platforms. On the other hand, S626-161 did not demonstrate neutralizing activity against Omicron variants beyond the BA.1 variant (Figure 1g). Although S626-161 had a lower neutralization potency against VOC than the other two, it was the only mAb which showed cross-reactivity to SARS-CoV-1 RBD and was able to neutralize bat coronaviruses WIV-1 and RsSHC014 (Figure 1f-g). These data suggest that S626-161 recognizes a conserved epitope that is shared between these sarbecovirus lineages, but is absent in BA.2. Additionally, compared to S728-1157 and S451-1140, S626-161 has a longer CDR-H3 which could provide an enhanced capability to recognize a highly conserved patch of residues shared across sarbecoviruses as described in a previous study^16^ (Figure S1). When comparing immunoglobulin heavy (IGHV) and light chain (IGLV or IGKV) variable genes of these three bnAbs with the available SARS-CoV-2 neutralizing mAbs database^12,17-25^, we found that heavy chain variable genes utilized by S728-1157 (IGHV3-66), S451-1140 (IGHV3-23) and S626-161 (IGHV4-39) have been previously reported to encode several potently neutralizing SARS-CoV-2 antibodies targeting the RBD^18,19,26,27^. However, only S728-1157 had unique heavy and light chain variable gene pairings that have not been reported in the database (Table S2), indicating that it is not public clonotype.

These three bnAbs (S451-1140, S626-161 and S728-1157) were characterized further to determine the binding breadth against SARS-CoV-2 VOCs (Figure 1h-k). The prefusion-stabilized spike containing two-proline substitutions in the S2 subunit (2P; diproline) has been shown to be a superior immunogen compared to the wildtype spike and is the basis of several current SARS-CoV-2 vaccines, including current mRNA-based vaccines^28,29^. More recently, spike protein stabilized with six prolines (6P; hexaproline) was shown to boost expression and be even more stable than the original diproline construct; as a result, it has been proposed for use in improving the next-generation of COVID-19 vaccines^30,31^. To determine if there are antigenicity differences between the diproline and hexaproline spike constructs, both immunogens were included in our test panel. As measured by ELISA assay, we found that three bnAbs bound 6P-WT spike antigen to a greater extent compared to WT-2P spike (Figure 1h-j). All three bnAbs showed comparable binding to the spikes of Alpha, Beta, Gamma and Delta viruses, relative to that of WT-2P (Figure 1h-j). However, the binding reactivity of these three bnAbs were substantially reduced against a panel of Omicron-family antigens (Figure 1h-k). S451-1140 binding was sensitive to mutations found in BA.1 and BA.2, resulting in largely decrease in binding and a 31-fold decrease in neutralization against these variants compared with WT-2P antigen and D614G virus, respectively (Figure 1g, i, k). The sarbecovirus-cross neutralizing mAb, S626-161 also showed 1.7 to 3.9-fold reduced binding to spike BA.1 antigens which may be affected in a 2-fold reduction in neutralization activity against BA.1 (Figure 1g, j, k). For the most potent bnAb, S728-1157, binding to Omicron antigens was substantially reduced by greater than 1.7-fold (range of 1.7-to 5.5-fold) compared with WT-2P spike but was unaffected in neutralizing activity (Figure 1g, h, k). Notably, all three bnAbs showed over 3-fold increased binding to spike BA.1-6P compared with the BA.1-2P version, suggesting a better accessibility of bnAbs to the hexaproline spike BA.1 construct. In addition to ELISA, biolayer interferometry (BLI) was used to quantify the binding rate and equilibrium constants (k_on_, k_off_, and K_D_) of these three bnAbs to a panel of spike antigens (Figure S2b-d). The recognition k_on_ rates of Fabs were 1.5 to 3.3-fold faster to hexaproline spikes, showing that the antibodies bound to the 6P construct more rapidly than to 2P. This is expected if the epitopes are more exposed on the RBD in the open state on the hexaproline spike (Figure S2c). Except for S626-161, off-rate of the Fabs were also longer such that the overall K_D_ showed that S728-1157 and S451-1140 bound to the hexaproline spike with substantially greater affinity (Figure S2c-d). The increased off rates further suggest partial occlusion of the binding site on diproline spike. The improved binding to hexaproline spike was even more notable for whole dimeric IgG by the 1:2 interaction model and for all three bnAbs, consistent with exposure of multiple epitopes with 6P stabilization allowing improved avidity (Figure S2b-d). Taken together, these results suggest that the epitopes targeted may be comparatively more accessible on the 6P-stabilized spike with the RBD in the open state. Structural analyses were next performed to verify this conjecture.

### Structural analysis of broadly neutralizing monoclonal antibodies

As a first approximation of epitopes bound, an ELISA competition assay was used to determine whether the three broadly-neutralizing mAbs shared any overlap with our current panel of mAbs and a collection of mAbs with known epitope specificities from previous studies^12,32,33^, plus two other mAbs currently in clinical use (LY-CoV555 (Eli Lilly)^34^ and REGN10933 (Regeneron)^35^). The binding sites of S451-1140 and S728-1157 partially overlapped with CC12.3^33,36^, a class 1 neutralizing antibody, and most class 2 antibodies, including LY-CoV555 and REGN10933, but not with class 3 and class 4 antibodies (Figure 2a). S626-161 shared a notable overlap in binding region with class 1 CC12.3, several class 4 antibodies including CR3022, and other unclassified antibodies, while having some partial overlap with several class 2 and one class 3 antibodies (Figure 2a). Analogously, competition BLI assay revealed that S451-1140 and S728-1157 strongly competed with one another for binding to spike WT-6P, whereas S626-161 did not (Figure S3). Overall, these data suggest S451-1140 and S728-1157 recognize similar epitopes that are distinct from S626-161.

**Figure 2:**
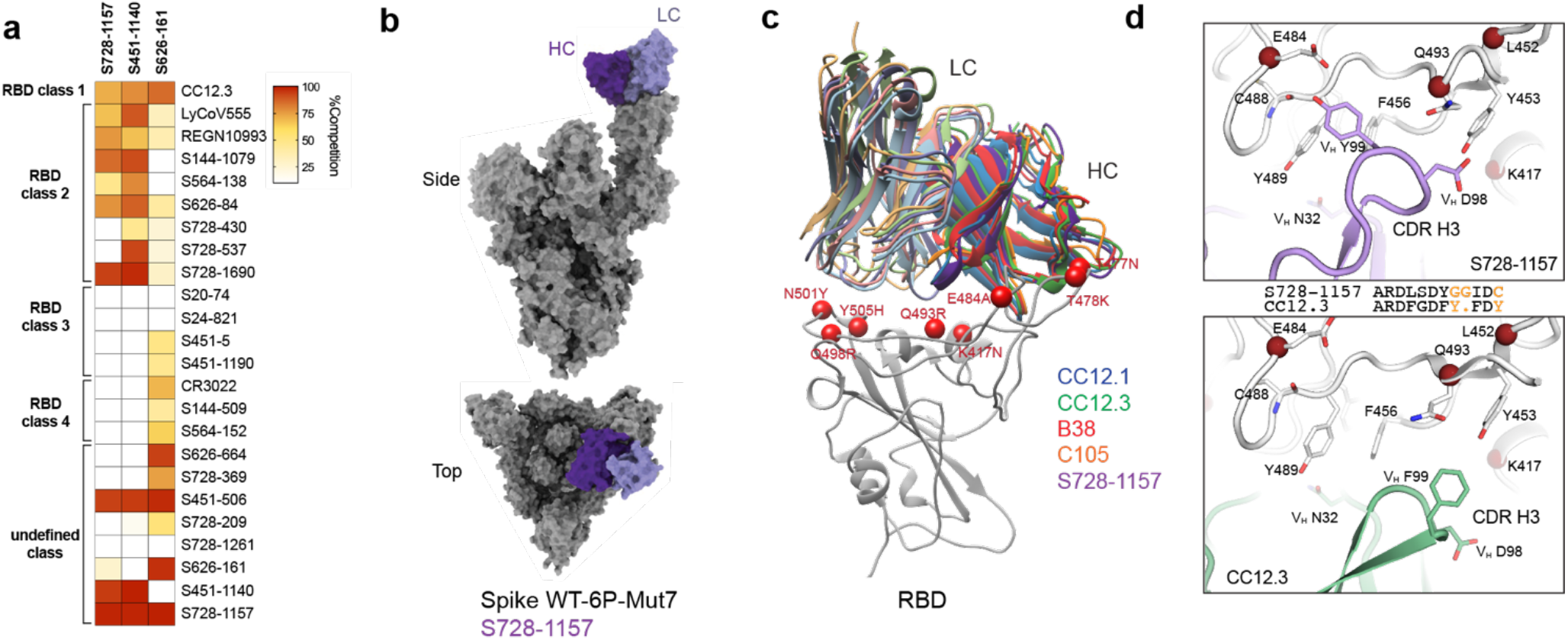
Mechanism of broad neutralization of S728-1157. **(a)** Epitope binning of broadly neutralizing RBD-reactive mAbs. Heatmap demonstrating the percentage of competition between each RBD-reactive mAb from previous studies^12,20,33-35^ with three broadly neutralizing mAbs, S728-1157, S451-1140 and S626-161. Data are representative of two independent experiments performed in triplicate. (**b)** Surface representation of the model derived from the cryoEM map of spike WT-6P-Mut7 in complex with IgG S728-1157. The heavy chain is shown in dark purple, light chain in light purple, and the spike protein in gray. Although we observe full mAb occupancy in the cryo-EM map, only one Fv is shown here. (**c)** Structural comparison of S728-1157 to other RBS-A antibodies such as CC12.1 (PDB ID: 6XC2, blue), CC12.3 (PDB ID: 6XC4, green), B38 (PDB ID: 7BZ5, red), and C105 (PDB ID: 6XCN, orange). The heavy chains are a darker shade, and the light chains are a lighter shade of their respective colors. Omicron BA.1 mutations near the epitope interface are shown as red spheres. (**d)** CDR-H3 forms distinct interactions with SARS-CoV-2 RBD between S728-1157 and CC12.3. Sequence alignment of CDR-H3 of the two antibodies are shown in the middle with non-conserved residues shown in orange.

S728-1157 was encoded by IGHV3-66 and possessed a short complementarity determining region 3 (CDR-H3). Notably, mAbs that bind the receptor binding site (RBS) in binding mode 1 (i.e. RBS-A or class 1 site), typified by CC12.1, CC12.3, B38, and C105^15,25,27,36-38^, tend to use IGHV3-53/3-66 and are sensitive to VOC mutations^39^. However, the CDR-H3 region of S728-1157 is highly distinct from other antibodies of this class, potentially accounting for its broader activity. To understand the structural basis of broad neutralization by S728-1157 at this epitope, we resolved a cryo-electron microscopy (cryo-EM) structure (Figure 2b) of IgG S728-1157 in complex with spike WT-6P-Mut7, a version of spike WT-6P possessing interprotomer disulfide bond at C705 and C883, at ∼3.3 Å global resolution (Figure S4e). Using symmetry expansion, focused classification, and refinement methods, we achieved local resolution at the RBD-Fv interface to ∼4Å (Figure S4e and Table S6). A crystal structure of S728-1157 Fab was determined at 3.1 Å resolution and used to build the atomic model at the RBD-Fv interface. Our structures confirm that S728-1157 binds the RBS-A (or class 1) epitope in the RBD-up conformation (Figure 2b and Figure S4e), similar to other IGHV3-53/3-66 antibodies (Figure 2c). Steric blockage of the angiotensin converting enzyme 2 (ACE2) binding site by S728-1157 explains its high neutralization potency against SARS-CoV-2. The _32_NY_33_ motif and _53_SGGS_56_ motif^36^ in S728-1157 CDR-H1 and-H2 interact with the RBD in almost the same way as CC12.3 (Figure S4b-c). However, V_H 98_DY_99_ in S728-1157 CDR-H3 forms more extensive interactions including both hydrophobic and polar interactions with the RBD, compared to V_H 98_DF_99_ in CC12.3 (Figure 2d and Table S5). The diglycine V_H 100_GG_101_ in S728-1157 CDR-H3 may also facilitate more extensive binding compared to V_H_ Y_100_ in CC12.3 likely due to the flexibility in the glycine residues that lead to a different conformation of the tip of the CDR-H3 loop and a relative shift of residues at _98_DY_99_.

Although the Omicron VOCs have extensive mutations in the RBD (Figure 2c and Figure S2a), most of these residues do not make interactions with or are dispensable for binding to S728-1157, as binding is still observed (Figure S4a). From our spike WT-6P-Mut7 + Fab S728-1157 model, Y505 to V_L_ Q31, and E484 to V_H_ Y99 are predicted to make hydrogen bonds (Figure S4d and Table S5), which have the potential to be disrupted by Omicron mutations Y505H and E484A. However, a Y505H mutation would still allow for a hydrogen bond with V_L_ Q31 and an E484A mutation would add another hydrophobic side chain near hydrophobic residues V_L_ Y99, F456, and Y489. These contacts may explain the mechanism which enabled S728-1157 to retain neutralizing activity (Figure 1g), albeit reduced binding reactivity against spike BA.1 antigen, which is in turn possibly due to the function of Omicron mutations in altering the conformational landscape of the spike protein^40^. Notably, while the variable genes were well-mutated, all but one of the contact residues between the CDR-H3 of S728-1157 and the VOC were predicted to be germline encoded and not introduced by somatic mutations, likely limiting the number of existing memory B cells of this class that could be further adapted by somatic mutation to protect against VOC strains (Figure S1, Table S5). Overall, our structural studies revealed the basis of broad neutralization of S728-1157 that can accommodate most mutations in the SARS-CoV-2 VOCs.

### S728-1157 reduces replication of SARS-CoV-2 Delta and Omicron variants in Syrian hamsters

To evaluate the protective efficacy of our broadly neutralizing mAbs, we utilized a golden Syrian hamster infection model that has been widely used for SARS-CoV-2 infection. Hamsters received 5 mg/kg of individual mAbs or an irrelevant antigen (ebolavirus glycoprotein)-specific isotype control via intraperitoneal injection one day post-infection with SARS-CoV-2 viruses. Lung and nasal tissues were collected at 4 days post-infection (dpi) (Figure 3a). Therapeutic administration of S728-1157 resulted in reduced titers of wildtype, BA.1 Omicron and Delta variants in both the nasal turbinates and lungs of infected hamsters (Figure 3b-d). Interestingly, the effect of S728-1157 in the lungs was dramatic, reducing wildtype viral loads by ∼10^4^ PFU, and BA.1 Omicron by ∼10^5^ PFU, with the viral titers of the latter being completely abolished (Figure 3c). In contrast to *in vitro* neutralization, S451-1140 did not reduce BA.1 Omicron viral replication in lung and nasal turbinates, indicating the disconnect between *in vitro* neutralization and *in vivo* protection for S451-1140 (Figure 3e). In comparison, S626-161 administration resulted in significant but marginal reductions in lung viral titers following wildtype and BA.1 challenge (Figure 3f-g). These data underscore that to precisely define broadly protective mAbs, evaluating protection efficacy in parallel with neutralization activity is required. Overall, S728-1157 represents a promising mAb with broad neutralization efficacy against SARS-CoV-2 variants that is capable of dramatically reducing wildtype, Delta and BA.1 replication *in vivo*.

**Figure 3:**
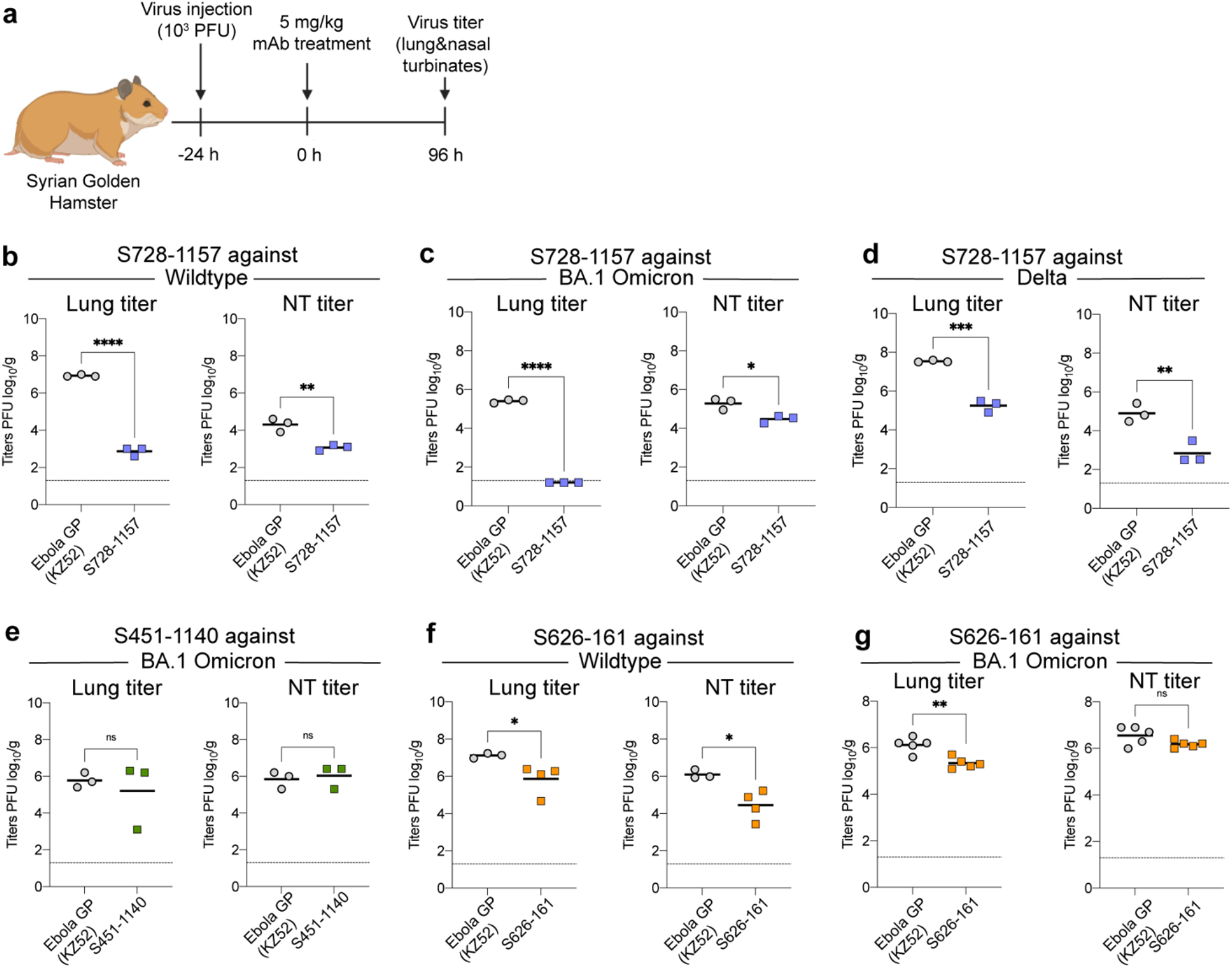
Protective efficacy of broadly neutralizing antibodies against SARS-CoV-2 infection in hamster. Schematic illustrating the in vivo experiment schedule **(a)**. Lung and nasal turbinate (NT) viral replication SARS-CoV-2 are shown for hamster treated therapeutically with **(b-d)** S728-1157 (n=3) **(e)** S451-1140 (n=3) and **(f-g)** S626-161 (n=4) at day 4 post-challenge with SARS-CoV-2 compared with control mAb, anti-Ebola surface glycoprotein (KZ52) antibody. Dashed horizontal lines represent the limit of detection (LOD) of the experiment. P-values in **(b-g)** were calculated using Unpaired t-test. The infected SARS-CoV-2 viruses are detailed in **Table S4**.

### SARS-CoV-2 infection rarely elicits potent S728-1157-like cross-neutralizing mAbs

Given the cross-neutralization and prophylactic potential of S728-1157, we sought to evaluate whether S728-1157-like antibodies are commonly induced among polyclonal responses in SARS-CoV-2 patients. To assess this, we performed competition ELISAs using convalescent serum to detect anti-RBD antibody titers that could compete for binding with S728-1157 (Figure 4a). Subjects were divided into three groups based on their magnitude of antibody responses, as defined previously^12,13^. Although high- and moderate-responders had higher titers of S728-1157-competitive serum antibodies compared to low-responders (Figure 4b), the titers were quite low across all groups suggesting that it is uncommon to acquire high levels of S728-1157-like antibodies in polyclonal serum following wildtype SARS-CoV-2 infection. In addition to S728-1157, we tested the competition of convalescent serum with other mAbs, including S451-1140 and S626-161, LY-CoV555, REGN10933, CR3022, and CC12.3. Similar to S728-1157, we observed relatively low titers of antibodies competing with S451-1140, S626-161, LY-CoV555, REGN10933 and CC12.3 in polyclonal serum from most of the convalescent individuals (Figure 4c-f, h). Nonetheless, high-responders tended to have significantly higher titers against those neutralizing mAbs than low-responders (Figure 4b-f, h). In contrast, antibodies targeting the CR3022 epitope site were more pronounced in convalescent individuals, suggesting the enrichment of class 4 RBD antibodies in polyclonal serum (Figure 4g). Notably, there was no difference in titers of CR3022 across the three responder groups, suggesting that CR3022-site antibodies were largely induced during wildtype SARS-CoV-2 infection. Interestingly, as compared to CC12.3, S728-1157 was detected at 4-fold lower levels in the serum of high-responders. Thus, despite class 1 antibodies being frequently induced by natural infection and vaccination^17,26,27,41-44^, our data suggest that S728-1157-like antibodies represent a subset of this class that are comparatively rare.

**Figure 4:**
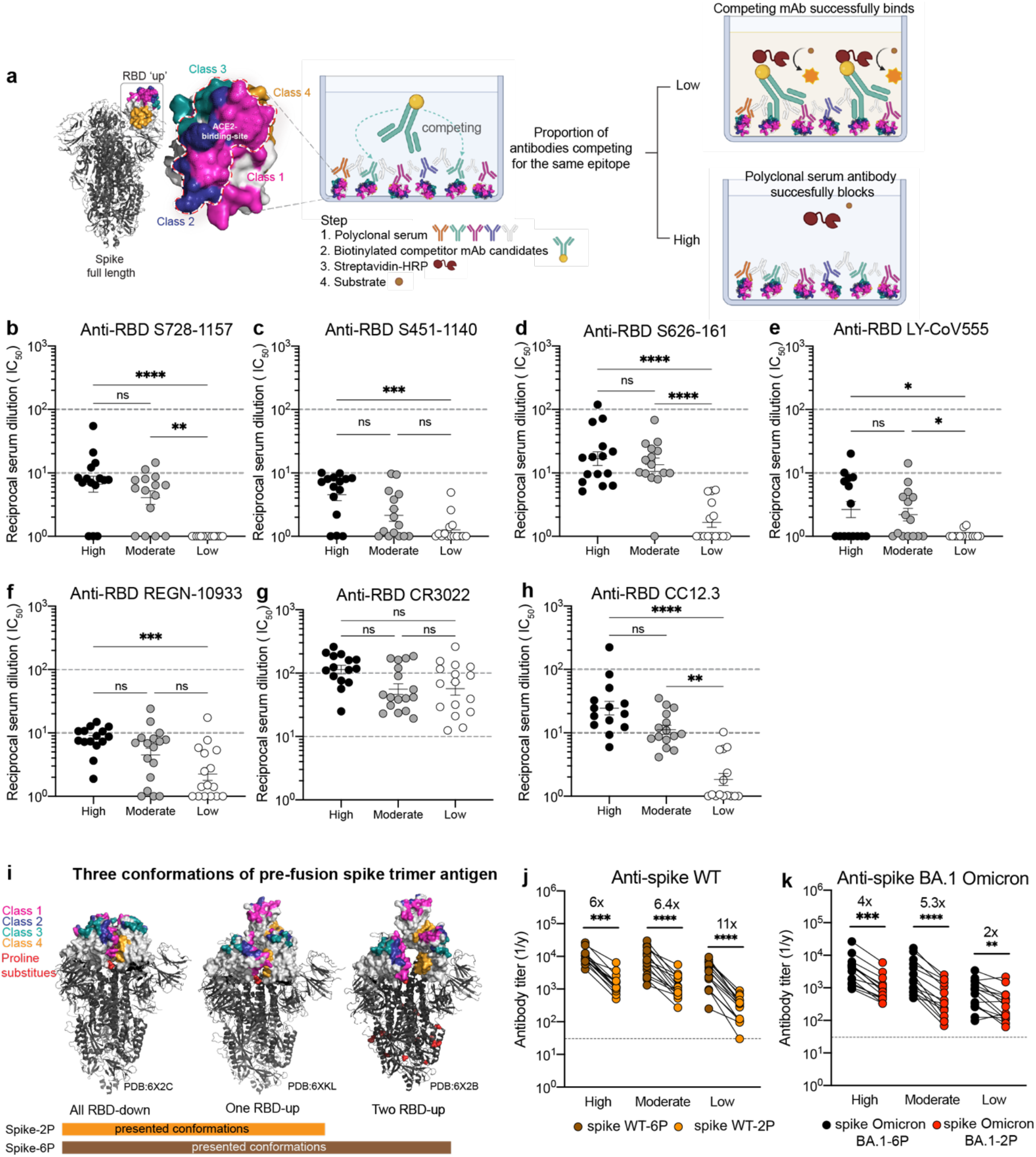
Convalescent serum antibody competition with broadly neutralizing RBD-reactive mAbs and comparison of serum antibody response against spike 6P-versus 2P-stabilized. Schematic diagram for experimental procedure of serum competitive ELISA **(a)**. Half-maximal inhibitory concentration (EC_50_) of polyclonal antibody serum from convalescent individuals that could compete with broadly neutralizing mAbs (competitor mAb): S728-1157 **(b)**, S451-1140 **(c)** and S626-161 **(d)**, therapeutic neutralizing mAbs LY-CoV555 **(e)**, REGN-10933 **(f)**, non-neutralizing mAb CR3022 **(g)** and well-defined class 1 mAb CC12.3 **(h)**. The reciprocal serum dilutions in **b-h** are showed as Log1P of the IC_50_ of serum dilution that can achieve 50% competition with the competitor mAb of interest. The statistical analysis in **b-h** was determined using Kruskal-Wallis with Dunn’s multiple comparison test. Representative three conformations of pre-fusion spike trimer antigen observed in the previous structural characterization of SARS-CoV-2 stabilized by 2P and 6P^31,47^ **(i)**. Endpoint titer of convalescent sera against SARS-CoV-2 spike wildtype (WT) **(j)** and Omicron BA.1 **(k)** in two versions of spike substituted by 2P and 6P. Data in **b-h and j-k** are representative of two independent experiments performed in duplicate. Wilcoxon matched-pairs signed rank test was used to compare the anti-spike antibody titer against 2P and 6P in **j-k**. Fold change indicated in **j-k** is defined as the mean fold change.

Lastly, we examined the difference in reactivity to 2P-versus 6P-stabilized spike in our convalescent cohort sera (Figure 4i-k). We found that all three responder groups mounted anti-spike reactive antibodies against 6P-stabilized spike wildtype to a greater extent than 2P-stabilized spike wildtype, by a factor of 6 to 11-fold (Figure 4j), indicating that the major antigenic epitopes were better exhibited or stabilized on 6P-stablized antigen. Using the same samples, high and moderate responders also had lower anti-spike antibodies against BA.1-2P than BA.1-6P, by 4 to 5-fold (Figure 4k). Of note, low responders had a smaller fold change in binding reactivity against spike BA.1 Omicron-2P and 6P (2-fold reduction) compared to wildtype-2P and 6P spike (11-fold reduction) (Figure 4j-k), suggesting that serum antibody against BA.1 Omicron-reactive epitopes may be limited in low responder subjects. Overall, these data suggest that there is improved polyclonal binding induced by natural infection to 6P-stabilized spike, both for wildtype and Omicron viruses.

## Discussion

In this study, we identify three potent bnAbs isolated from memory B cells of individuals who had recovered from SARS-CoV-2 infection during the initial wave of the COVID-19 pandemic. Among them, S728-1157 maintains substantial binding reactivity and had consistent neutralizing activity against all tested SARS-CoV-2 VOC including Omicron BA.1, BA.2, BA.2.75, BL.1 (BA.2.75+R346T), BA.4 and BA.5, and was able to substantially reduce infectious viral titers following Delta and BA.1 infection in hamsters. We found convalescent serum from our cohort contained low concentrations of antibodies that compete with S728-1157 (a class 1 antibody) and class 2 epitope mAbs. This suggests that S728-1157 is somewhat unique from other antibodies targeting class 1 epitopes and is infrequently induced in the RBD-specific memory B cells pool. Instead, in our cohort natural infection preferably induced antibodies targeting the CR3022 (class 4) epitope; antibodies of this specificity are also often non-neutralizing or less potently neutralizing than RBS-targeting antibodies. These data are complementary to our previous findings that demonstrated that an abundance of class 3 antibodies in convalescent sera may contribute to neutralizing activity against Alpha and Gamma variants, whereas a lack of class 2 antibodies may account for reduced neutralization capability against Delta^12^. Notwithstanding, the breadth of activity against Omicron variants of most of these RBS-targeting antibodies (RBS-A/class 1, RBS-B,C/class 2 and RBS-D/class 3) is reported to be highly limited^10,39,45^. This is consistent with experimental evidence documenting that convalescent unvaccinated patients showed a marked reduction of neutralizing activity against Omicron BA.1^9^. This phenomenon highlights the need to shape the antibody repertoire toward broadly conserved, protective epitopes, as typified by S728-1157.

The structures herein illustrated that S728-1157 bound the RBS-A/class 1 epitope in the ‘up’ conformation RBD. This epitope is more readily accessible on 6P-stabilized spikes, which present two RBDs in the ‘up’ state, as compared to 2P spikes which presents only one^28,31,46,47^. The S728-1157 was isolated after natural infection; in such contexts, the odds of inducing S728-1157-like clones are likely higher given that the RBD must be able to adopt an up conformation, even transiently, to bind to ACE2, thereby exposing this epitope. Using extensive interactions between CDR-H3 and the RBD, S728-1157 can accommodate key mutations in VOC spikes, indicating this antibody is unlike majority of IGHV3-53/3-66 RBS-A/class 1 antibodies^27,48,49,50^. S728-1157 also uses a different light chain (IGLV3-9) compared to other non-broad antibodies such as CC12.3 (IGKV3-20), which may affect overall binding conformation; however, our analysis indicates that there is limited direct interaction between the S728-1157 light chain and the RBD. Most of the CDR-H3 contact residues critical for VOC cross-reactivity in this interaction are germline-encoded and not introduced by somatic mutations, suggesting that most memory B cells encoding IGHV3-53/66 class antibodies could not acquire this degree of cross-reactivity by further affinity maturation, which is a consideration for vaccines tailored to help induce such antibodies. While it may be challenging to design vaccines that can specifically elicit S728-1157-like antibodies with select CDR-H3s capable of overcoming the VOC mutations, it is encouraging that IGHV-gene restriction is observed in other potent SARS-CoV-2 neutralizing mAbs studies^12,17-25^. Ultimately, such a vaccination approach may be feasible through iterative immunization with optimized RBD immunogens, as has been previously reported in the influenza literature^51-55^.

Although many mutations have been observed in the class 1 antigenic site^15^, with regards to the S728-1157 epitope 13/15 total RBD contact residues, and 2/3 CDR-H3-bound RBD contact residues, are conserved within Omicron and all other VOCs. This suggests that the RBD region where the S728-1157 epitope is found may include residues critical for its dynamic function and viral fitness and would therefore be less tolerant of mutations and antigenic drift than surrounding class 1 site residues. If this is the case, the tendency for this particular epitope to be lost as viral variants evolve should be reduced, making characterization of S728-1157 and similar antibodies and epitopes important for variant-resistant vaccines or mAb therapeutic development.

In summary, our study identifies broadly neutralizing antibodies that may inform immunogen design for next-generation variant-proof coronavirus vaccines or serve as mAb therapeutics that are resistant to SARS-CoV-2 evolution. In particular, in terms of combined potency and breadth, S728-1157 appears to be the best-in-class antibody isolated to date. Given that this antibody is predicted to be preferentially induced by 6P-stabilized recombinant spike proteins or whole virus, these findings suggest that hexaproline modification could benefit future vaccine constructs to optimally protect against future SARS-CoV-2 variants and other sarbecoviruses.

## Acknowledgements

We thank the study participants for their willingness to enroll in this study. We are grateful for the clinical staff at the University of Chicago Medicine Plasma Transfusion Program for their assistance in collecting the sample and transfer to the lab. We also kindly thank the University of Chicago CAT Facility (RRID SCR_017760) and the University of Chicago Genomics Facility (RRID SCR_019196) for assisting in sorting and sequencing samples. We thank Henry Tien for technical support with the crystallization robot, and Robyn Stanfield for assistance in data collection. We thank Jeffrey Copps for producing the spike proteins used for electron microscopy. We thank Bill Anderson and Charles Bowman for maintaining the microscope facility and for technical assistance. We are grateful to the staff of the Stanford Synchrotron Radiation Lightsource (SSRL) beamline 12-1 for assistance. Use of resources of the SSRL, SLAC National Accelerator Laboratory is supported by the U.S. Department of Energy, Office of Science, Office of Basic Energy Sciences under Contract No. DE-AC02–76SF00515. Extraordinary facility operations were supported in part by the DOE Office of Science through the National Virtual Biotechnology Laboratory, a consortium of DOE national laboratories focused on the response to COVID-19, with funding provided by the Coronavirus CARES Act. The SSRL Structural Molecular Biology Program is supported by the DOE Office of Biological and Environmental Research, and by the National Institutes of Health, National Institute of General Medical Sciences (including P41GM103393).

## Author contributions

Conceptualization, SC, PCW; Methodology, SC, PJH, HL, JLT, GO, JT, MH, SAE, YF, DW; Sample Collection, NYZ, HLT; Resources, GS, FK, DNS, IAW, ABW, YK. Investigation, SC, PJH, HL, JLT, JJM, LL, MK, TM; Writing – Original Draft, SC, HL, JLT; Writing – Review and Editing, all authors; Funding, PCW, FK, DNS, IAW, ABW, YK. Supervision, PCW, IAW, ABW, YK.

## Supplementary Figures and Tables

**Figure S1:**
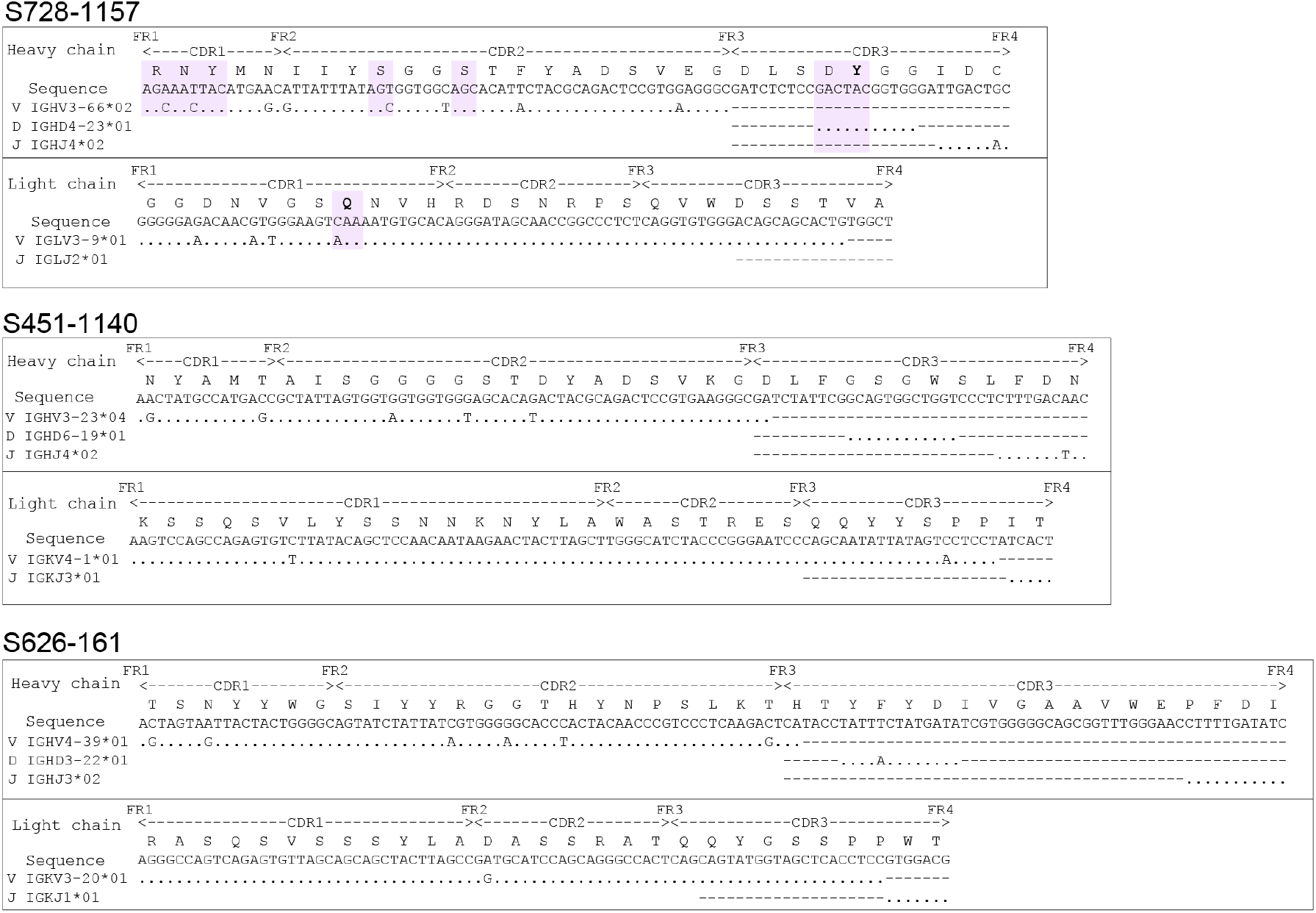
Amino acid and nucleotide sequences of complementarity-determining region (CDR) of heavy chain and light chain of the three bnAbs. Contacting residues within CDR of S728-1157 and SARS-CoV-2 are highlighted as light purple. Genetic information for each antibody is in **Table S2**.

**Figure S2:**
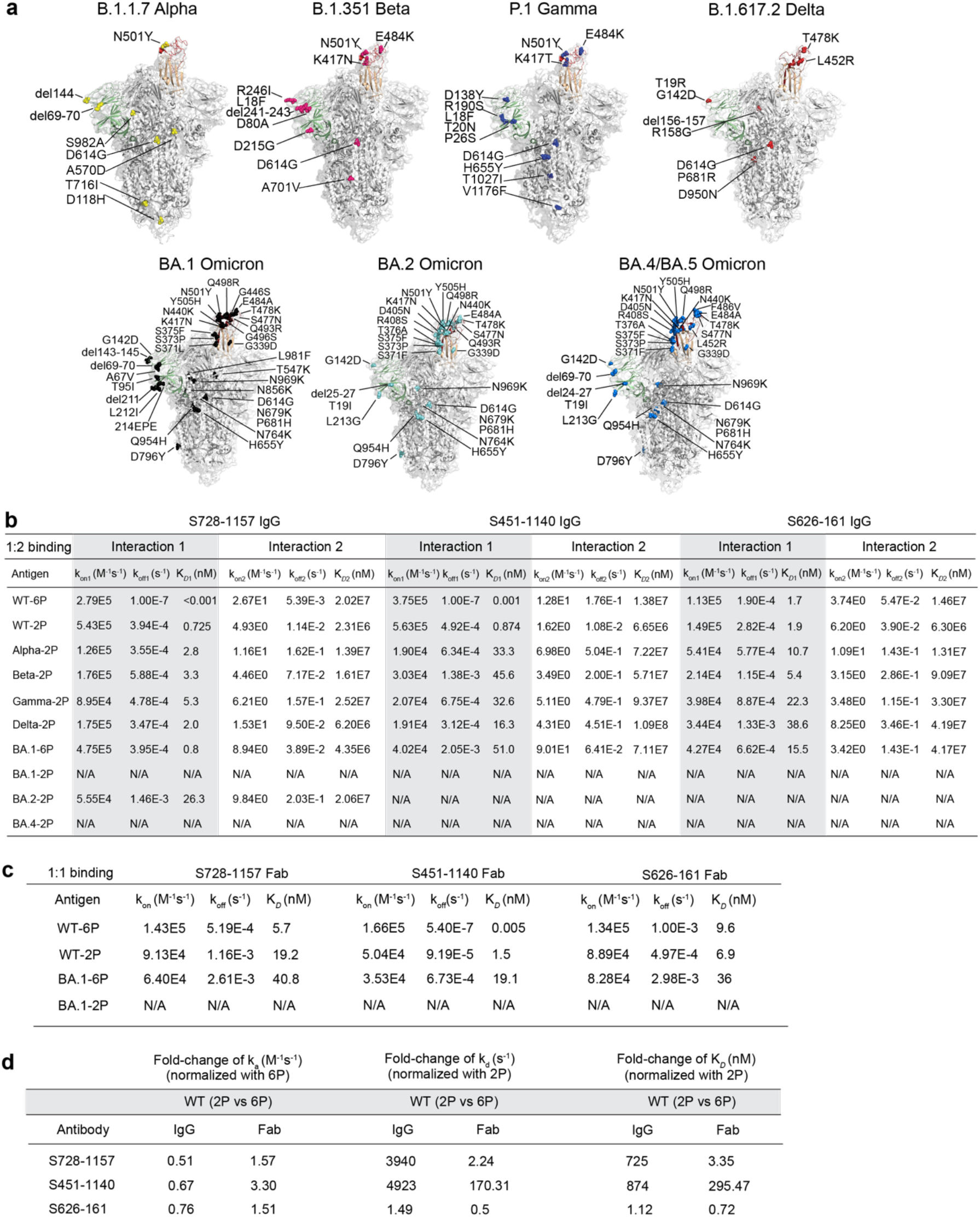
Broadly neutralizing RBD-reactive mAbs activity against SARS-CoV-2 and emerging variants. **a**, Structural models for the full-length spike protein variants and amino acid substitutions that encoded in B.1.1.7 Alpha, B.1.351 Beta, P.1 Gamma, B.1.617.2 Delta and Omicron, BA.1, BA.2 and BA.4. The structural models in **a** are modified from PDB ID: 6XM4. **b**, The table illustrating the binding rate and equilibrium constants (k_on_, k_off_, and affinity binding K_D_) measured by BLI of S728-1157, S451-1140 and S626-161 IgG in response to the panel of SARS-CoV-2 VOCs (either former or current VOCs). **c**, The binding rate comparison of Fabs of S728-1157, S451-1140 and S626-161 in responding to spike WT and BA.1-6P and 2P constructs. The binding traces of IgG and Fab analyzed by BLI were represented by the 1:2 and 1:1 interaction model, respectively. **d**, The fold-change of binding rate (K_on_, K_off_) and binding affinity (K_D_) between spike WT-6P and spike WT-2P bound by neutralizing RBD-reactive mAbs, whole IgG form and Fab. Data in **c-d** are representative of two independent experiments, the data from experiments that have the best fit (R^2^ > 0.90) are selected for analysis.

**Figure S3:**
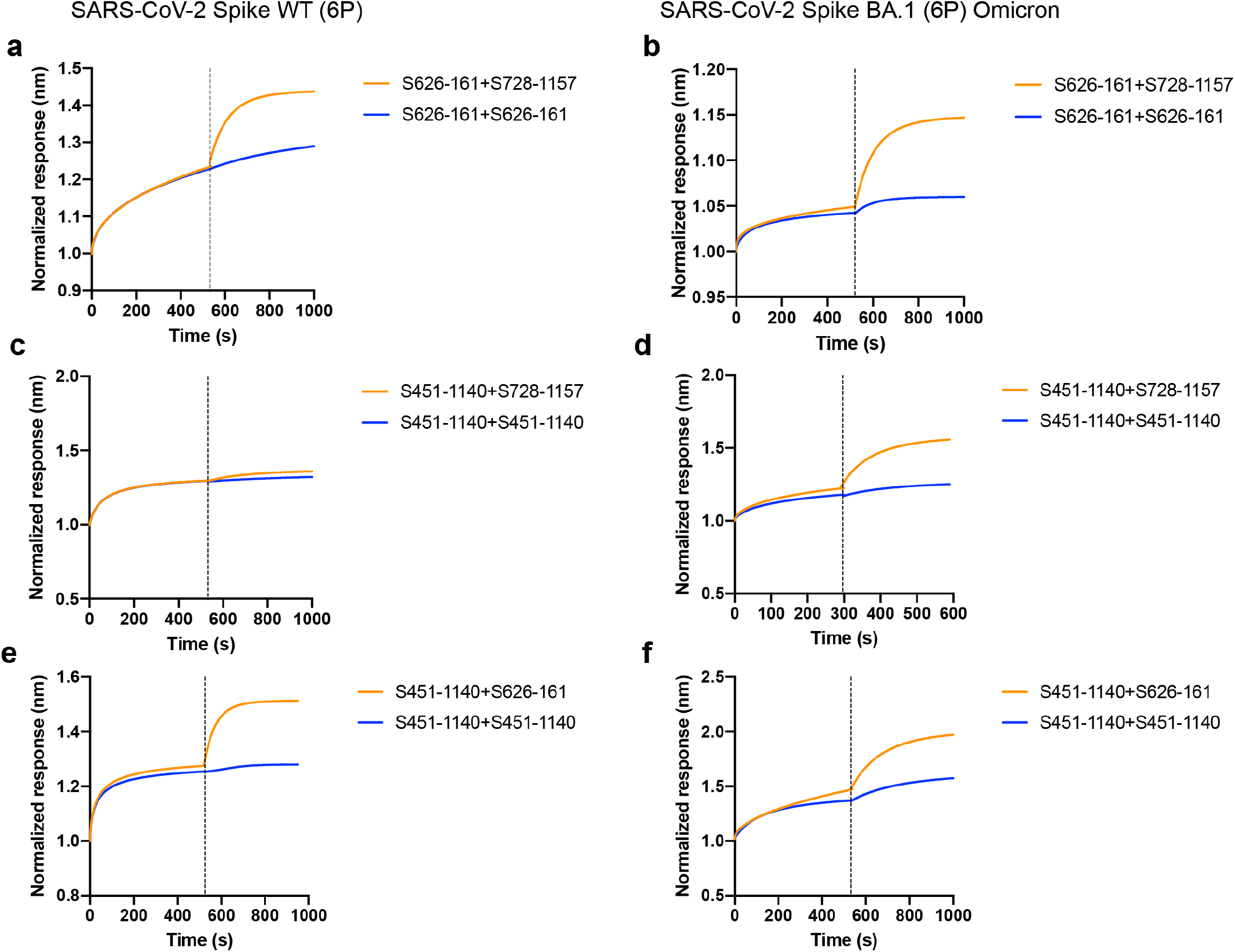
Biolayer interferometry analysis demonstrates binding affinity curves of three broadly neutralizing mAbs competing with each other in response to biotinylated spike wildtype (WT)-6P (left panel) and spike BA.1 Omicron-6P (right panel). **a-b**, S626-161 was firstly bound, followed by S728-1157 mAb as competing mAb. **c-d**, S451-1140 was firstly bound and competed with S728-1157 and **e-f**, S626-161. The response curve was normalized in relation to its starting response value.

**Figure S4.**
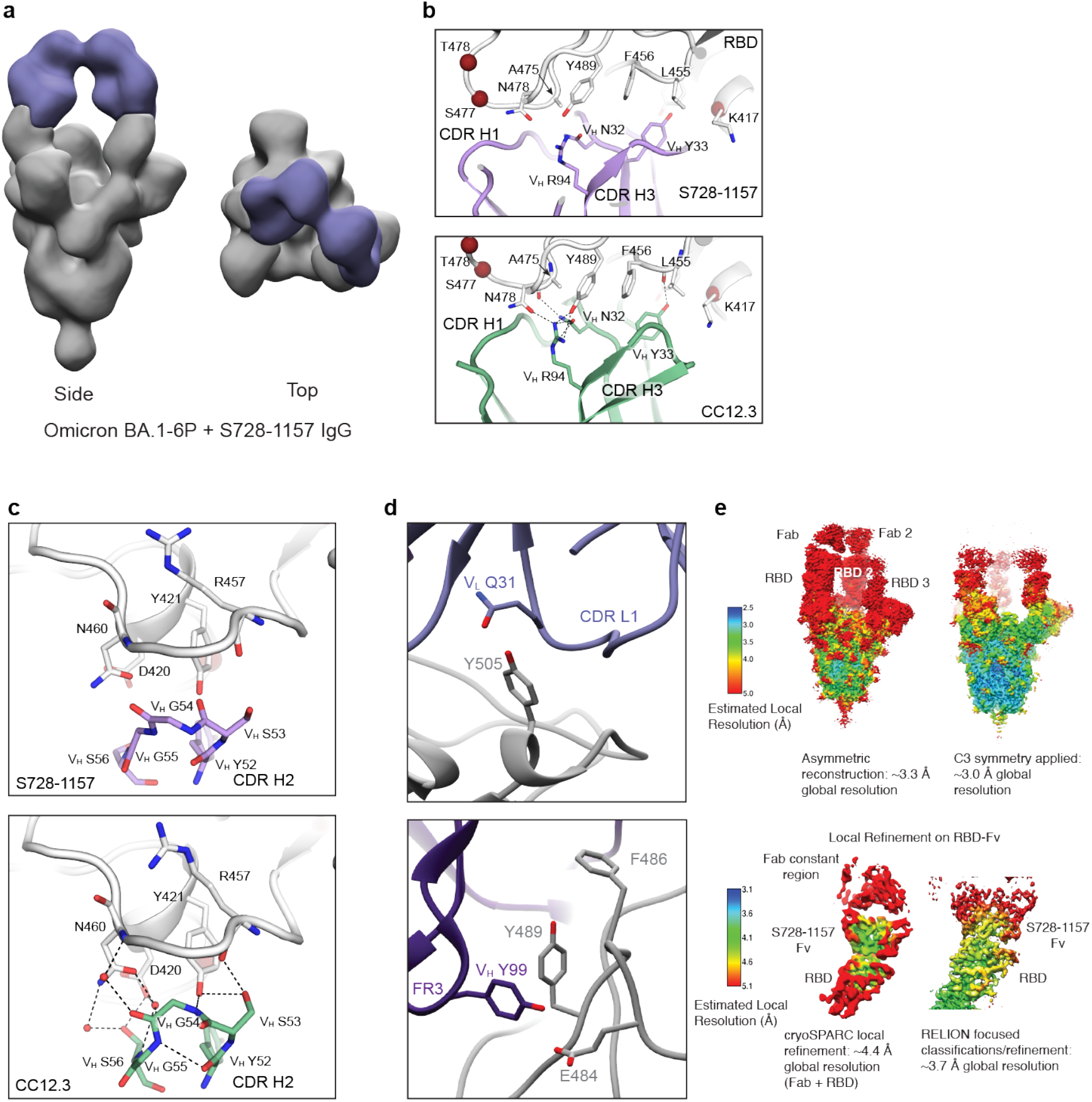
Structural analysis of S728-1157 binding to SARS-CoV-2 spike. (**a)** Three-dimensional (3D) reconstruction of Omicron BA.1-6P in complex with IgG S728-1157 shows binding by negative stain electron microscopy. The binding mode is the same as binding to spike WT-6P-Mut7 shown in **Figure 2b**. (**b)** CDR-H1 of S728-1157 forms similar interactions with SARS-CoV-2 RBD compared to another IGHV3-53 antibody CC12.3 (PDB ID: 6XC4). (**c)** CDR-H2 of S728-1157 forms similar interactions with the RBD compared to CC12.3 (PDB ID: 6XC4). (**d)** For spike WT-6P-Mut7 in complex with S728-1157, residues Y505 and V_L_ Q31, and E484 and V_L_ Y99 are predicted to make hydrogen bonds. Hydrophobic residues Y486 and Y489 are shown as well. Since S728-1157 binds spike Omicron BA.1-6P in the same way as to spike WT-6P-Mut7, it may accommodate the E484A and Y505H mutations in Omicron. (**e)** Local resolution estimates of the cryo-EM map (upper panel) and local refinement on the RBD-Fv after symmetry expansion using RELION (lower panel).

**Table S1:**
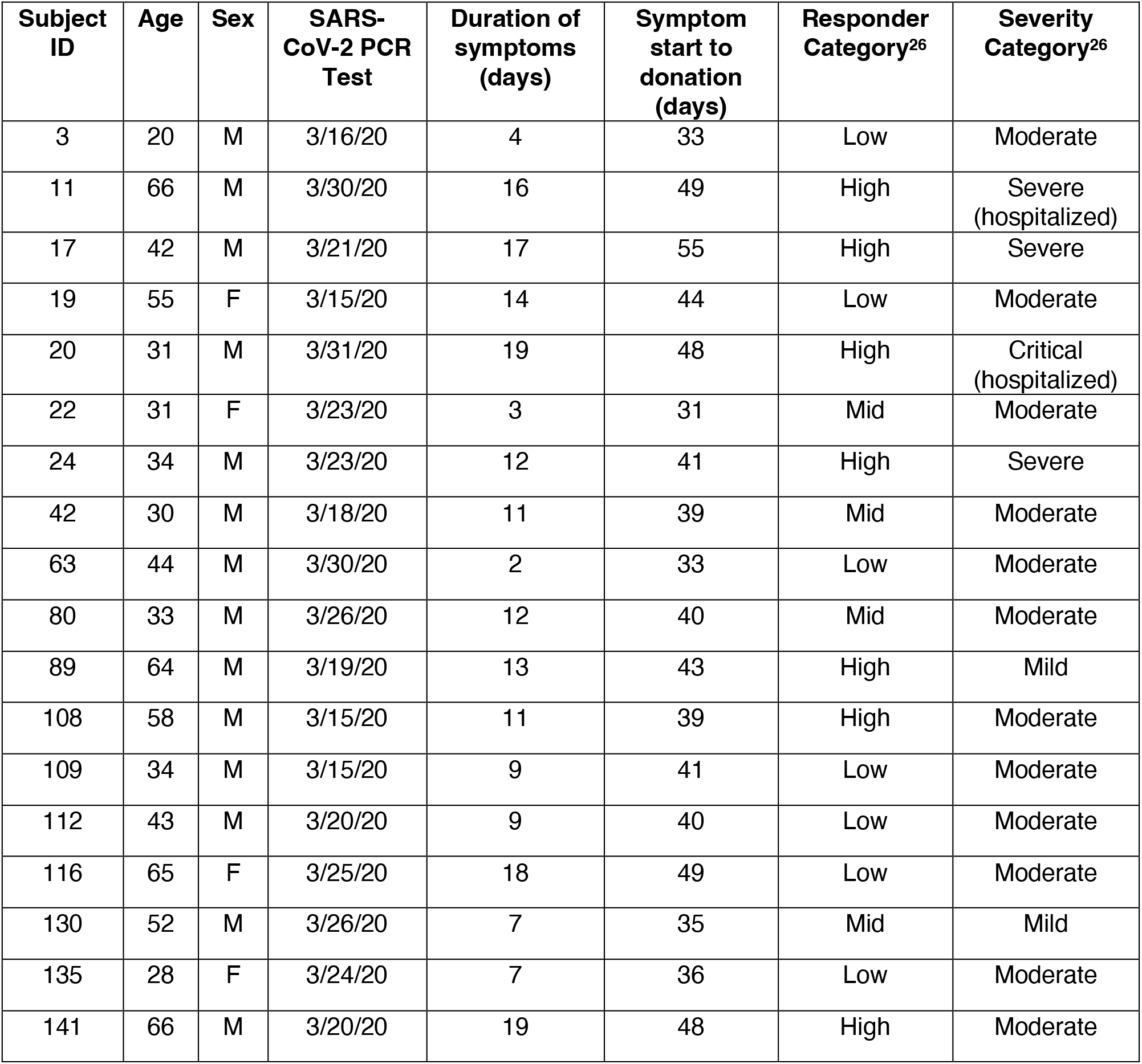
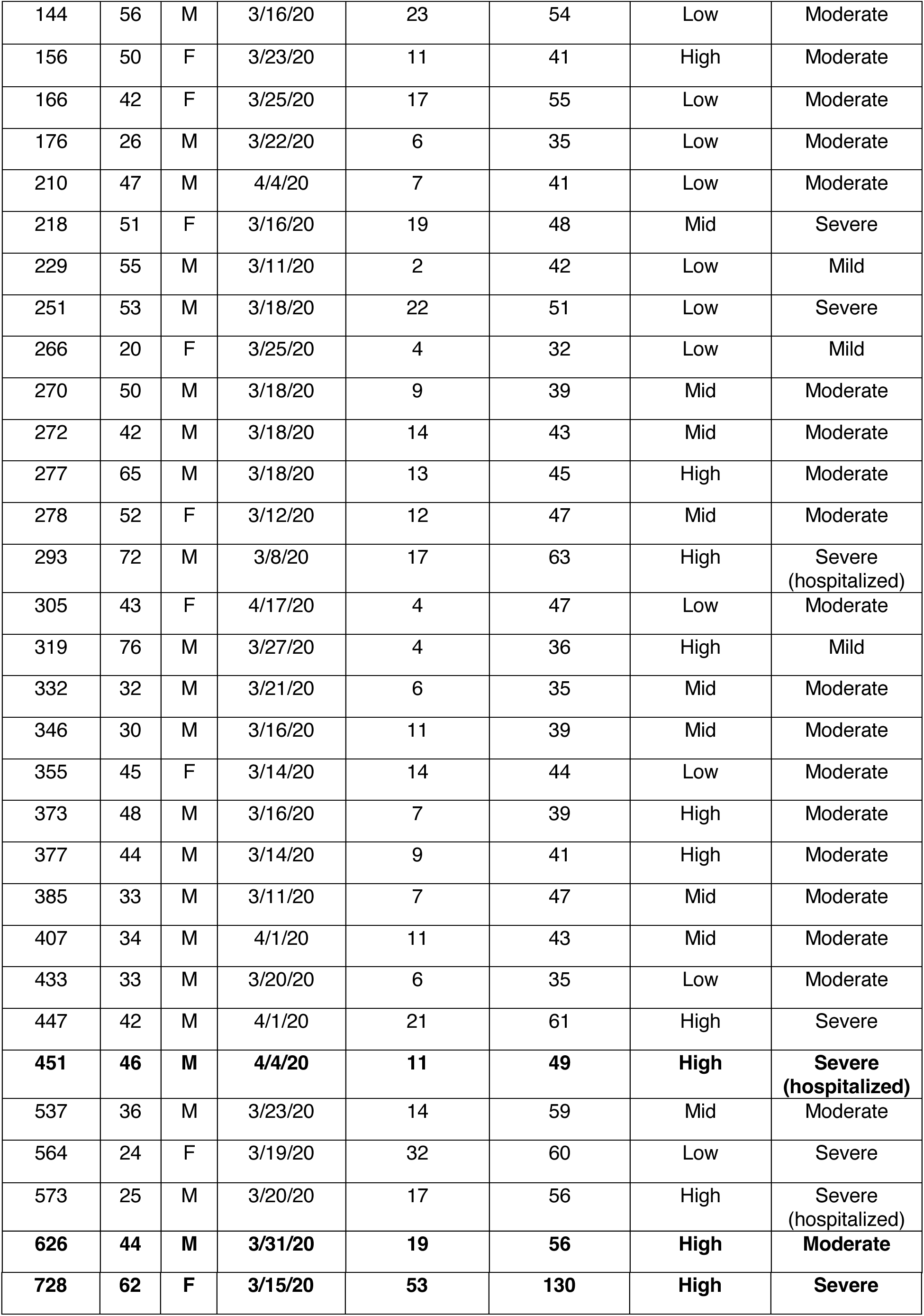
COVID-19 convalescent subjects. Related to Figure 1 and Figure 4. The mAbs from high responder subjects, S451, S626, S728 were characterized in this study. Responder group and severity were categorized in a previous study^13^. Serum antibody from each responder group were tested for competition ELISA with broad neutralizing mAbs, other therapeutic mAbs and non-neutralizing mAb.

**Table S2:**
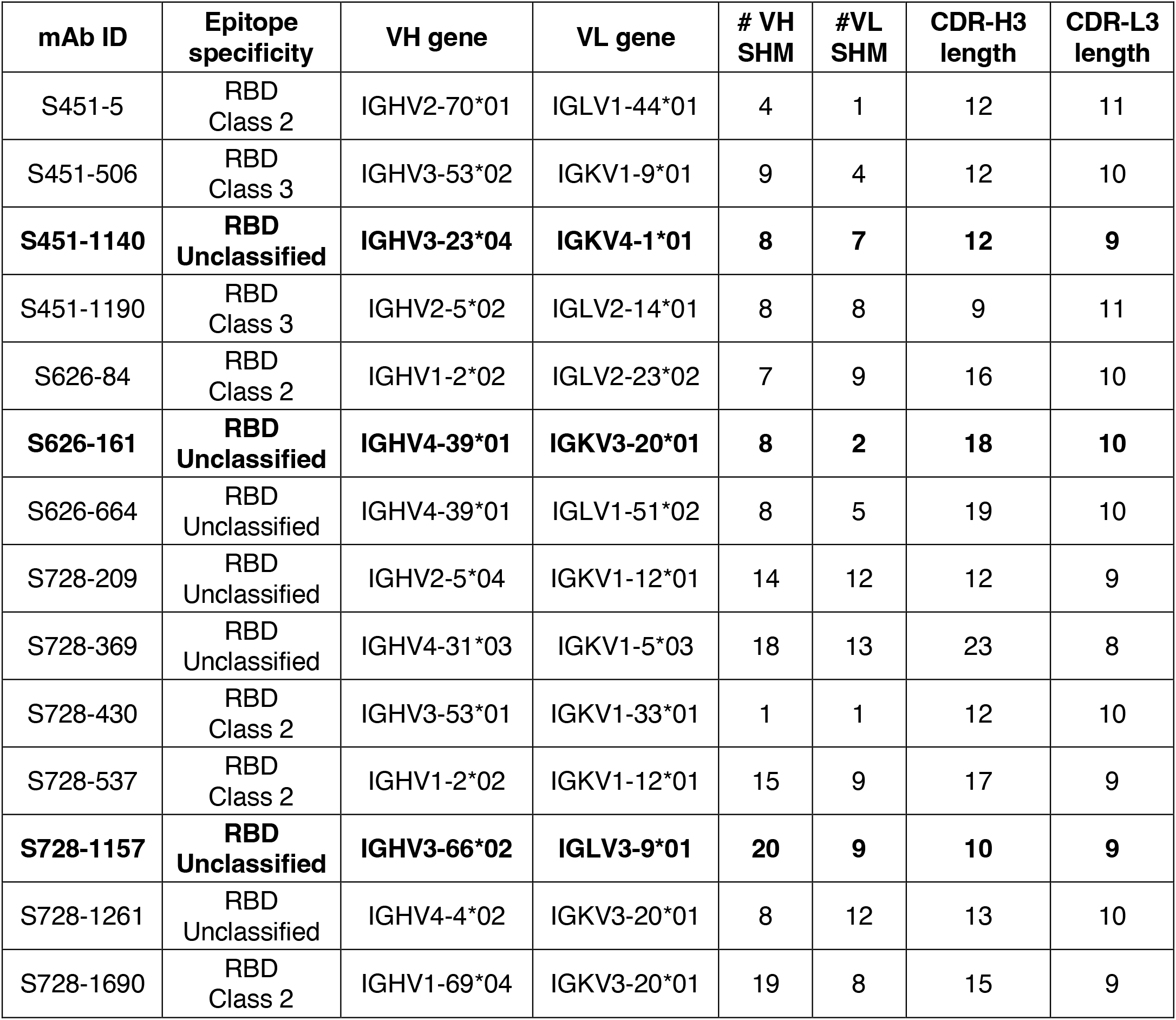
Characteristics of SARS-CoV-2 RBD-reactive mAbs. Related to Figure 1. Cross-neutralizing mAbs against D614G and B.1.351 Beta, B.1.,617.2 Delta, B.1.617.1 Kappa, B.1.621 Mu, BA.1 Omicron are bolded.

| mAb ID | Epitope specificity | VH gene | VL gene | # VH SHM | #VL SHM | CDR-H3 length | CDR-L3 length |
| --- | --- | --- | --- | --- | --- | --- | --- |
| S451-5 | RBD Class 2 | IGHV2-70*01 | IGLV1-44*01 | 4 | 1 | 12 | 11 |
| S451-506 | RBD Class 3 | IGHV3-53*02 | IGKV1-9*01 | 9 | 4 | 12 | 10 |
| **S451-1140** | **RBD Unclassified** | **IGHV3-23\*04** | **IGKV4-1\*01** | **8** | **7** | **12** | **9** |
| S451-1190 | RBD Class 3 | IGHV2-5*02 | IGLV2-14*01 | 8 | 8 | 9 | 11 |
| S626-84 | RBD Class 2 | IGHV1-2*02 | IGLV2-23*02 | 7 | 9 | 16 | 10 |
| **S626-161** | **RBD Unclassified** | **IGHV4-39\*01** | **IGKV3-20\*01** | **8** | **2** | **18** | **10** |
| S626-664 | RBD Unclassified | IGHV4-39*01 | IGLV1-51*02 | 8 | 5 | 19 | 10 |
| S728-209 | RBD Unclassified | IGHV2-5*04 | IGKV1-12*01 | 14 | 12 | 12 | 9 |
| S728-369 | RBD Unclassified | IGHV4-31*03 | IGKV1-5*03 | 18 | 13 | 23 | 8 |
| S728-430 | RBD Class 2 | IGHV3-53*01 | IGKV1-33*01 | 1 | 1 | 12 | 10 |
| S728-537 | RBD Class 2 | IGHV1-2*02 | IGKV1-12*01 | 15 | 9 | 17 | 9 |
| **S728-1157** | **RBD Unclassified** | **IGHV3-66\*02** | **IGLV3-9\*01** | **20** | **9** | **10** | **9** |
| S728-1261 | RBD Unclassified | IGHV4-4*02 | IGKV3-20*01 | 8 | 12 | 13 | 10 |
| S728-1690 | RBD Class 2 | IGHV1-69*04 | IGKV3-20*01 | 19 | 8 | 15 | 9 |

**Table S3:** Antigen information and resource. Proline substitutions are indicated as italic. **Related to Figure 1 and Figure S2**.

| Antigen | S1 NTD | RBD | S1 CTD | S2 | Source |
| --- | --- | --- | --- | --- | --- |
| Spike FL, trimer |  |  |  |  |  |
| Wildtype(WT)-2P | - | - | - | *K986P*,<br>*V987P* | Krammer lab |
| Wildtype(WT)-6P | - | - | - | *F817P*,<br>*A829P*,<br>*A899P*,<br>*A942P*,<br>*K986P*,<br>*V987P* | Krammer lab |
| B.1.1.7 Alpha-2P | del69-70,<br>del144 | N501Y | A570D,<br>D614G,<br>P681H | T716I,<br>S982A,<br>*K986P*,<br>*V987P*,<br>D1118H | Sather lab |
| B.1.351 Beta-2P | L18F,<br>D80A, D215G,<br>del241-243,<br>R246I | K417N,<br>E484K,<br>N501Y | D614G | A701V,<br>*K986P*,<br>*V987P* | Sather lab |
| P.1 Gamma-2P | L18F,<br>T20N,<br>P26S, D138Y,<br>R190S | K417T,<br>E484K,<br>N501Y | D614G,<br>H655Y | *K986P*,<br>*V987P*,<br>T1027I,<br>V1176F | Sather lab |
| B.1.617.2 Delta-2P | T19R, G142D,<br>del156-157,<br>R158G | L452R,<br>T478K, | D614G,<br>P681R | D950N,<br>*K986P*,<br>*V987P* | Sather lab |
| BA.1 Omicron-2P | A67V,<br>H69del,<br>V70del,<br>T95I,<br>G142D,<br>V143del,<br>Y144del,<br>Y145del,<br>N211del,<br>L212I,<br>insert214EPE | G339D,<br>S371L,<br>S373P,<br>S375F,<br>K417N,<br>N440K,<br>G446S,<br>S477N,<br>T478K,<br>E484A,<br>Q493R,<br>G496S,<br>Q498R,<br>N501Y,<br>Y505H | T547K,<br>D614G,<br>H655Y,<br>N679K,<br>P681H | N764K,<br>D796Y,<br>N856K,<br>Q954H,<br>N969K,<br>L981F,<br>*K986P*,<br>*V987P* | Sather lab |
| BA.1 Omicron-6P | A67V,<br>H69del,<br>V70del,<br>T95I,<br>G142D,<br>V143del,<br>Y144del, | G339D,<br>S371L,<br>S373P,<br>S375F,<br>K417N,<br>N440K,<br>G446S, | T547K,<br>D614G,<br>H655Y,<br>N679K,<br>P681H | *V705C*,<br>N764K,<br>D796Y,<br>*F817P*,<br>*A829P*,<br>N856K,<br>*T883C*, | Ward lab |
|  | Y145del,<br>N211del,<br>insert214EPE | S477N,<br>T478K,<br>E484A,<br>Q493R,<br>G496S,<br>Q498R,<br>N501Y,<br>Y505H |  | A899P,<br>A942P,<br>Q954H,<br>N969K,<br>L981F,<br>K986P,<br>V987P |  |
| BA.2 Omicron-2P | T19I,<br>L24del,<br>P25del,<br>P26del,<br>A27S,<br>G142D,<br>V213G, | G339D,<br>S371F,<br>S373P,<br>S375F,<br>T376A,<br>D405N,<br>R408S,<br>K417N,<br>N440K,<br>S477N,<br>T478K,<br>E484A,<br>Q493R,<br>Q498R,<br>N501Y,<br>Y505H | D614G,<br>H655Y,<br>N679K,<br>P681H, | N764K,<br>D796Y,<br>Q954H,<br>N969K,<br>K986P,<br>V987P | Sather lab |
| BA.4 Omicron-2P | T19I,<br>L24del,<br>P25del,<br>P26del,<br>A27S,<br>H69del,<br>V70del,<br>G142D,<br>V213G, | G339D,<br>S371F,<br>S373P,<br>S375F,<br>T376A,<br>D405N,<br>R408S,<br>K417N,<br>N440K,<br>L452R,<br>S477N,<br>T478K,<br>E484A,<br>F486V,<br>Q498R,<br>N501Y,<br>Y505H, | D614G,<br>H655Y,<br>N679K,<br>P681H, | N764K,<br>D796Y,<br>Q954H,<br>N969K,<br>K986P,<br>V987P | Sather lab |
| RBD |  |  |  |  |  |
| WT | - | - | - | - | In-house |
| R346S | - | R346S | - | - | In-house |
| K417N | - | K417N | - | - | In-house |
| K417V | - | K417V | - | - | Krammer lab |
| K417T | - | K417T | - | - | In-house |
| G446V | - | G446V | - | - | In-house |
| N439K | - | N439K | - | - | Krammer lab |
| L452R | - | L452R | - | - | In house |
| S477N | - | S477N | - | - | In-house |
| E484K | - | E484K | - | - | Krammer lab |
| F486A | - | F486A | - | - | In-house |
| F486Y | - | F486Y | - | - | In-house |
| N487Q | - | N487Q | - | - | In-house |
| Y489F | - | Y489F | - | - | In-house |
| Q493A | - | Q493A | - | - | In-house |
| Q493N | - | Q493N | - | - | In-house |
| N501Y | - | N501Y | - | - | In-house |
| Y505A | - | Y505A | - | - | In-house |
| Y505F | - | Y505F | - | - | In-house |
| K417N/E484K/<br>L452R/N501Y | - | K417N/<br>E484K/<br>L452R/<br>N501Y | - | - | In-house |
| SARS-CoV-1 RBD WT |  |  |  |  | In-house |
| MERS-CoV RBD WT |  |  |  |  | In-house |

**Table S4:** SARS-CoV-2 virus information and resource. Related to Figure 1 and 3.

| Virus | S1 NTD | RBD | S1 CTD | S2 | Source |
| --- | --- | --- | --- | --- | --- |
| D614G | - | - | D614G | - | 2019-nCoV/USA-WA1/2020 D614G |
| B.1.351<br>Beta | L18F,<br>D80A,<br>D215G,<br>L241del,<br>L242del,<br>A243del | K417N,<br>E484K,<br>N501Y | D614G | A701V | hCoV-19/USA/MD-HP01542/2021 |
| P.1<br>Gamma | L18F,<br>T20N,<br>P26S,<br>D138Y,<br>G181V,<br>R190S | K417T,<br>E484K,<br>N501Y | D614G,<br>H655Y | T1027I,<br>V1176F | hCoV-19/Japan/TY7-501/2021 from BEI |
| B.1.621<br>Mu | in3T,<br>T95I,<br>Y144S,<br>Y145N, | R346K,<br>E484K,<br>N501Y | D614G,<br>P681H | D950N | hCoV-19/USA/WI-UW-4340/2021 |
| B.1.617.1<br>Kappa | G142D,<br>E154K | L452R,<br>E484Q | D614G,<br>P681R | Q1071H,<br>H1101D | hCoV-19/USA/CA-Stanford-15_S02/2021 from BEI |
| B.1.617.2<br>Delta | T19R,<br>T95I,<br>G142D,<br>E156G,<br>F157del,<br>R158del | L452R,<br>T478K | D614G,<br>P681R | D950N | hCoV-19/USA/WI-UW-5250/2021 |
| BA.1<br>Omicron | A67V,<br>H69del,<br>V70del,<br>T95I,<br>G142D,<br>V143del,<br>Y144del, | G339D,<br>S371L,<br>S373P,<br>S375F,<br>K417N,<br>N440K,<br>G446S, | T547K,<br>D614G,<br>H655Y,<br>N679K,<br>P681H | N764K,<br>D796Y,<br>N856K,<br>Q954H,<br>N969K,<br>L981F | hCoV-19/USA/WI-WSLH-221686/2021 |
|  | Y145del,<br>N211del,<br>L212I,<br>ins214EPE | S477N,<br>T478K,<br>E484A,<br>Q493R,<br>G496S,<br>Q498R,<br>N501Y,<br>Y505H |  |  |  |
| BA.2<br>Omicron | T19I,<br>delL24,<br>delP25,<br>delP26,<br>A27S,<br>G142D,<br>V213G | G339D,<br>S371F,<br>S373P,<br>S375F,<br>T376A,<br>D405N,<br>R408S,<br>K417N,<br>N440K,<br>S477N,<br>T478K,<br>E484A,<br>Q493R,<br>Q498R,<br>N501Y,<br>Y505H | D614G,<br>H655Y,<br>N679K,<br>P681H | N764K,<br>D796Y,<br>Q954H,<br>N969K | hCoV-19/Japan/UT-NCD1288-2N/2022 |
| BA.2.75<br>Omicron | T19I,<br>delL24,<br>delP25,<br>delP26,<br>A27S,<br>G142D,<br>K147E,<br>W152R,<br>F157L,<br>I210V,<br>V213G,<br>G257S | G339H,<br>S371F,<br>S373P,<br>S375F,<br>T376A,<br>D405N,<br>R408S,<br>K417N,<br>N440K,<br>G446S,<br>N460K,<br>S477N,<br>T478K,<br>E484A,<br>Q498R,<br>N501Y,<br>Y505H | D614G,<br>H655Y,<br>N679K,<br>P681H | N764K,<br>D796Y,<br>Q954H,<br>N969K | hCoV-19/Japan/TY41-716/2022 |
| BA.4<br>Omicron | T19I,<br>delL24,<br>delP25,<br>delP26,<br>A27S,<br>delH69,<br>delV70,<br>G142D,<br>V213G | G339D,<br>S371F,<br>S373P,<br>S375F,<br>T376A,<br>D405N,<br>R408S,<br>K417N,<br>N440K,<br>L452R,<br>S477N,<br>T478K, | D614G,<br>H655Y,<br>N679K,<br>P681H, | N764K,<br>D796Y,<br>Q954H,<br>N969K | hCoV-19/USA/MD-HP30386-PIDNBNVCCQ/2022 |
|  |  | F486V,<br>E484A,<br>Q498R,<br>N501Y,<br>Y505H, |  |  |  |
| BA.5<br>Omicron | T19I,<br>delL24,<br>delP25,<br>delP26,<br>A27S,<br>delH69,<br>delV70,<br>G142D,<br>V213G | G339D,<br>S371F,<br>S373P,<br>S375F,<br>T376A,<br>D405N,<br>R408S,<br>K417N,<br>N440K,<br>L452R,<br>S477N,<br>T478K,<br>F486V,<br>E484A,<br>Q498R,<br>N501Y,<br>Y505H, | D614G,<br>H655Y,<br>N679K,<br>P681H, | N764K,<br>D796Y,<br>Q954H,<br>N969K | SARS-CoV-2/human/USA/COR-22-063113/2022 |
| BL.1<br>Omicron | T19I,<br>delL24,<br>delP25,<br>delP26,<br>A27S,<br>G142D,<br>K147E,<br>W152R,<br>F157L,<br>I210V,<br>V213G,<br>G257S | G339H,<br>R346T,<br>S371F,<br>S373P,<br>S375F,<br>T376A,<br>D405N,<br>R408S,<br>K417N,<br>N440K,<br>G446S,<br>N460K,<br>S477N,<br>T478K,<br>E484A,<br>Q498R,<br>N501Y,<br>Y505H | D574V,<br>D614G,<br>H655Y,<br>N679K,<br>P681H | N764K,<br>D796Y,<br>Q954H,<br>N969K | SARS-CoV-2/human/USA/WI-UW-12980/2022 |

**Table S5:** Pairs of S728-1157 and spike-WT-6P-Mut7 residues within predicted hydrogen bonding distances. Calculated using EpitopeAnalyzer^63^ using a cutoff distance of 3.4 Å. **Related to Figure 2 and Figure S4**.

| # | RBD<br>Residue<br>[Atom] | Ab<br>Residue<br>[atom] | Antibody<br>Region | Distance<br>(Å) | RBD residue<br>mutated in<br>Omicron VOC | RBD residue<br>conserved across<br>all VOC's |
| --- | --- | --- | --- | --- | --- | --- |
| 1 | T415 [OG] | S56 [OG1] | CDRH2 | 2.78 | No | Yes |
| 2 | Y421 [OH] | S53 [O] | CDRH2 | 2.73 | No | Yes |
| 3 | Y453 [OH] | D98 [OD1] | CDRH3 | 3.5 | No | Yes |
| 4 | L455 [O] | Y33 [OH] | CDRH1 | 3.29 | No | Yes |
| 5 | R457 [O] | S53 [OG] | CDRH2 | 3.25 | No | Yes |
| 6 | Y473 [OH] | R31 [O] | CDRH1 | 2.76 | No | Yes |
| 7 | Y473 [OH] | S53 [OG] | CDRH2 | 3.26 | No | Yes |
| 8 | Q474 [O] | R31 [NH1] | CDRH1 | 3.08 | No | Yes |
| 9 | A475 [O] | L28 [N] | CDRH1 | 3.05 | No | Yes |
| 10 | A475 [O] | N32 [ND2] | CDRH1 | 2.98 | No | Yes |
| 11 | E484 [OE2] | Y99 [OH] | CDRH3 | 2.61 | Yes | No |
| 12 | N487 [ND2] | G26 [O] | FR1 | 3.01 | No | Yes |
| 13 | C488 [O] | Y99 [OH] | CDRH3 | 3.25 | No | Yes |
| 14 | Y489 [OH] | R94 [NH1] | FR3 | 2.64 | No | Yes |
| 15 | Y505 [OH] | Q31 [NE2] | CDRL1 | 2.62 | Yes | No |

**Table S6.** Cryo-EM data collection, refinement and model building statistics. Related to Figure 2 and Figure S4.

| Map | S728-1157 + SARS-CoV-2-6P-Mut7 (global refinement) | S728-1157 + SARS-CoV-2-6P-Mut7 (focused refinement) |
| --- | --- | --- |
| EMDB | EMD-27112 | EMD-27113 |
| Data collection |  |  |
| Microscope | Thermo Fisher Titan Krios |  |
| Voltage (kV) | 300 |  |
| Detector | Gatan K2 Summit |  |
| Recording mode | Counting |  |
| Nominal magnification | 130kx |  |
| Movie micrograph pixelsize (Å) | 1.045 |  |
| Dose rate (e<sup>-</sup>/[(camera pixel)*s]) | 6.017 |  |
| Number of frames per movie micrograph | 36 |  |
| Frame exposure time (ms) | 250 |  |
| Movie micrograph exposure time (s) | 9 |  |
| Total dose (e<sup>-</sup>/Å<sup>2</sup>) | 50.0 |  |
| Defocus range (μm) | -0.8 to -1.5 |  |
| EM data processing |  |  |
| Number of movie micrographs | 1,718 | 1,718 |
| Number of molecular projection images in map | 151,948 | 29,595 |
| Symmetry | C1 | C1 |
| Map resolution (FSC 0.143; Å) | 3.3 | 3.7 |
| Map sharpening B-factor ($\text{\AA}^2$) | -85.3 | -71.1 |
| **Structure Building and Validation** |  |  |
| *Number of atoms in deposited model* |  |  |
| SARS-CoV-2-6P-Mut7 | *n/a* | 20,759 |
| Fab Fv | *n/a* | 1,653 |
| Glycans | *n/a* | 182 |
| MolProbity score | *n/a* | 1.07 |
| Clashscore | *n/a* | 1.66 |
| Map correlation coefficient | *n/a* | 0.75 |
| EMRinger score | *n/a* | 2.57 |
| d FSC model (0.5; $\text{\AA}$) | *n/a* | 3.8 |
| *RMSD from ideal* |  |  |
| Bond length ($\text{\AA}$) | *n/a* | 0.021 |
| Bond angles ($^\circ$) | *n/a* | 1.81 |
| *Ramachandran plot* |  |  |
| Favored (%) | *n/a* | 97.13 |
| Allowed (%) | *n/a* | 2.87 |
| Outliers (%) | *n/a* | 0.00 |
| Side chain rotamer outliers (%) | *n/a* | 0.08 |
| Cβ outliers (%) | *n/a* | 0.00 |
| PDB | *n/a* | 8d0z |

## Materials and Methods

### Monoclonal antibody isolation

We isolated a panel of RBD-reactive mAbs from peripheral blood mononuclear cells (PBMCs) of convalescent donors who previously had experienced symptomatic infection with SARS-CoV-2 (Table S1). The samples were collected during the first wave of the pandemic in May 2020, before other SARS-CoV-2 variants emerged. All studies were performed with the approval of the University of Chicago institutional review board (IRB20-0523). All participants provided prior written informed consent for the use of blood in research applications. This clinical trial was registered at ClinicalTrials.gov under identifier NCT04340050.

PBMCs were isolated from leukoreduction filters and frozen as described previously^21^. B cells were enriched from PBMCs via fluorescence-activated cell sorting (FACS). Cells were stained with CD19, CD3, and antigen probes conjugated oligo-fluorophore; cells of interest were identified as CD3^-^CD19^+^Antigen^+^. All mAbs were generated from oligo-tagged antigen bait-sorted cells identified through single-cell RNA sequencing (RNA-seq), as described previously^12,21^.

Antigen-specific B cells were selected to generate mAbs based on antigen-probe intensity analyzed by JMP Pro 15. Antibody heavy and light chain genes were synthesized and cloned into human IgG1 and human kappa or lambda light chain expression vectors by Gibson assembly as previously described^56^. The heavy and light chains of the corresponding mAb were transiently co-transfected into HEK293T cells. After transfection for 18 h, the transfected cells were supplemented with Protein-Free Hybridoma Medium Supernatant (PFHM-II, Gibco). The supernatant containing secreted mAb was harvested at day 4 and purified using protein A-agarose beads (Thermo Fisher) as detailed previously^56^.

### Recombinant spike protein expression

The recombinant D614G SARS-CoV-2 full-length (FL) spike, WT RBD, single RBD mutants (R346S, K417N, K417T, G446V, L452R, S477N, F486A, F486Y, N487Q, Y489F, Q493A, Q493N, N501Y, Y505A, Y505F), combination RBD mutant (K417N/E484K/L452R/NN501Y), SARS-CoV-1 RBD and MERS-CoV RBD were generated in-house. Briefly, the recombinant antigens were expressed using Expi293F cells. The gene of interest was cloned into mammalian expression vector (in-house modified AbVec) and transfected using ExpiFectamine 293 kit according to the manufacturer’s protocol. The supernatant was harvested at day 4 after transfection and incubated with Ni-nitrilotriacetic acid (Ni-NTA) agarose (Qiagen). The purification was carried out using gravity flow column and eluted with imidazole-containing buffer as previously described^57,58^. The eluate was buffering-exchanged with PBS using Amicon centrifugal unit (Millipore). The recombinant FL spikes derived from variants B.1.1.7 Alpha, B.1.351 Beta, P.1 Gamma, B.1.617.2 Delta, BA.1, BA.2 and BA.4 Omicron were produced in the Sather Laboratory at Seattle Children’s Research Institute. The K417V, N439K, E484K RBDs and recombinant FL spike WT-2P and 6P were produced in Krammer laboratory at the Icahn School of Medicine at Mount Sinai. The SARS-CoV-2-6P-Mut7 and spike BA.1 Omicron-6P were designed and produced as described in a previous study^59^. The protein sequences and resources for each antigen are listed in Table S3.

### Enzyme-linked immunosorbent assay (ELISA)

Recombinant SARS-CoV-2 spike/RBD proteins were coated onto high protein-binding microtiter plates (Costar) at 2 μg/ml in phosphate buffered saline (PBS) at 50 μl/well, and kept overnight at 4°C. Plates were washed with PBS containing 0.05% Tween 20 (PBS-T) and blocked with 150 μl of PBS containing 20% fetal bovine serum (FBS) for 1 h at 37°C. Monoclonal antibodies were serially diluted 3-fold starting from 10 μg/ml in PBS and incubated in the wells for 1 h at 37°C. Plates were then washed and incubated with horseradish peroxidase (HRP)-conjugated goat anti-human IgG antibody (Jackson ImmunoResearch, 1:1000) for 1 h at 37°C. After washing, 100 μl of Super AquaBlue ELISA substrate (eBioscience) was added per well. Absorbance was measured at 405nm on a microplate spectrophotometer (Bio-Rad). The assays were standardized using control antibodies with known binding characteristics in every plate, and the plates were developed until the absorbance of the control reached an optical density (OD) of 3.0. All mAbs were tested in duplicate, and each experiment was performed twice.

### Serum ELISA

High protein-binding microtiter plates were coated with recombinant SARS-CoV-2 spike antigens at 2 μg/ml in PBS overnight at 4°C. Plates were washed with PBS 0.05% Tween and blocked with 200 μl PBS 0.1% Tween + 3% skim milk powder for 1 hour at room temperature (RT). Plasma samples were heat-inactivated for 1 hour at 56°C before perform serology experiment. Plasma were serially diluted 2-fold in PBS 0.1% Tween + 1% skim milk powder. Plates were incubated with serum dilutions for 2 hours at RT. The HRP-conjugated goat anti-human Ig secondary antibody diluted at 1:3000 with PBS 0.1% Tween + 1% skim milk powder was used to detect binding of antibodies. After 1-hour of incubation, plates were developed with 100 μl SigmaFast OPD solution (Sigma-Aldrich) for 10 minutes. Then, 50 μl 3M HCl was used to stop the development reaction. Absorbance was measured at 490 nm on a microplate spectrophotometer (BioRad). End point titers were extrapolated from sigmoidal 4PL (where X is log concentration) standard curve for each sample. Limit of detection (LOD) is defined as the mean plus 3 S.D. of the O.D. signal recorded using plasma from pre-SARS-CoV-2 subjects. All calculations were performed in GraphPad Prism software (version 9.0).

### Competition ELISA

To determine the target epitope classification of RBD-reactive mAbs, competition ELISAs were performed using other mAbs with known epitope binding characteristics as competitor mAbs. Competitor mAbs were biotinylated using EZ-Link sulfo-NHS-biotin (Thermo Scientific) for 2h at room temperature (RT). The excess biotin of biotinylated mAbs was removed with 7k molecular weight-cutoff (MWCO) Zeba spin desalting columns (Thermo Scientific). Plates were coated with 2 μg/ml RBD antigen overnight at 4°C. Plates were blocked with PBS–20% FBS for 2h at RT, and the 2-fold dilution of the mAbs of an undetermined class, or serum, was added, starting at 20 μg/ml of mAbs and a 1:10 dilution of serum. After antibody incubation for 2h at RT, the biotinylated competitor mAb was added at a concentration twice that of its dissociation constant (K_D_) and incubated for another 2 h at RT together with the mAb or serum that was previously added. Plates were washed and incubated with 100 μl HRP-conjugated streptavidin (Southern Biotech) at a dilution of 1:1000 for 1 h at 37°C. The plates were developed with the Super AquaBlue ELISA substrate (eBioscience). To normalize the assays, the competitor biotinylated mAb was added in a well without any competing mAbs or serum as a control. Data were recorded when the absorbance of the control well reached and OD of 1.0-1.5. The percent competition between mAbs was then calculated by dividing a sample’s observed OD by the OD reached by the positive control, subtracting this value from 1, and multiplying by 100. For serum, ODs were log_10_-transformed and analyzed by nonlinear regression to determine the 50% inhibition concentration (IC_50_) values using GraphPad Prism software (version 9.0). The data were transformed to Log1P and plotted into graph representative of reciprocal serum dilution of the IC_50_ of serum dilution that can achieve 50% competition with the competitor mAb of interest. All mAbs were tested in duplicate, each experiment was performed two times independently, and values from two independent experiments were averaged.

### Plaque assays

Plaque assays were performed with SARS-CoV-2 variant viruses on Vero E6/TMPRSS2 cells (Table S4). Cells were cultured to achieve 90% confluency prior to being trypsinized and seeded at a density of 3×10^4^ cells/well in 96-well plates. On the following day, 10^2^ plaque-forming unit (PFU) of SARS-CoV-2 variant was incubated with 2-fold-diluted mAbs for 1h. The antibody-virus mixture was incubated with Vero E6/TMPRSS2 cells for 3 days at 37°C. Plates were fixed with 20% methanol and then stained with crystal violet solution. The complete inhibitory concentrations (IC_99_) were calculated using the log(inhibitor) versus normalized response (variable slope), performed in GraphPad Prism (version 9.0). All mAbs were tested in duplicate, and each experiment was performed twice.

### Focus reduction neutralization test (FRNT)

Focus reduction neutralization test (FRNT) were used to determine neutralization activities as an additional platform beside plaque assay. Serial dilutions of serum starting at a final concentration of 1:20 will be mixed with 10^3^ focus-forming units of virus per well and incubated for 1 h at 37 °C. A pooled pre-pandemic serum sample is served as a control. The antibody-virus mixture will be inoculated onto Vero E6/TMPRSS2 cells in 96-well plates and incubated for 1 h at 37 °C. An equal volume of methylcellulose solution was added to each well. The cells were incubated for 16 h at 37 °C and then fixed with formalin. After the formalin was removed, the cells were immunostained with a mouse monoclonal antibody against SARS-CoV-1/2 nucleoprotein [clone 1C7C7 (Sigma-Aldrich)], followed by a HRP-labeled goat anti-mouse immunoglobulin (SeraCare Life Sciences). The infected cells were stained with TrueBlue Substrate (SeraCare Life Sciences) and then washed with distilled water. After cell drying, the focus numbers were quantified by using an ImmunoSpot S6 Analyzer, ImmunoCapture software, and BioSpot software (Cellular Technology). The IC_50_ was calculated from the interpolated value from the log(inhibitor) versus normalized response, using variable slope (four parameters) nonlinear regression performed in GraphPad Prism (version 9.0).

### Negative stain electron microscopy

Spike BA.1 Omicron-6P was complexed with a 0.5-fold molar excess of IgG S728-1157 and incubated for 30 mins at room temperature. The complex was diluted to 0.03 mg/ml and deposited on a glow-discharged carbon-coated copper mesh grid. 2% uranyl formate (w/v) was used to stain the sample for 90 seconds. The negative stain dataset was collected on a Thermo Fisher Tecnai T12 Spirit (120keV, 56,000x magnification, 2.06 apix) paired with a FEI Eagle 4k x 4k CCD camera. Leginon^60^ was used to automate the data collection and raw micrographs were store in the Appion database^61^. Dogpicker^62^ picked particles and the dataset was processed in RELION 3.0^62^. UCSF Chimera^63^ was used for map segmentation and figure making.

### Cryo-electron microscopy and model building

SARS-CoV-2-6P-Mut7 was complexed with a 0.5-fold molar excess of IgG S728-1157 and incubated for 30 mins at room temperature. Grids were prepared using a Thermo Fisher Vitrobot Mark IV set to 4°C and 100% humidity. The complex, at 0.7 mg/ml, was briefly incubated with lauryl maltose neopentyl glycol (final concentration of 0.005 mM; Anatrace), deposited on a glow-discharged Quantifoil 1.2/1.3-400 mesh grid, and blotted for 3 seconds. The grid was loaded into a Thermo Fisher Titan Krios (130,000x magnification, 300 kEV, 1.045-A° pixel size) paired with a Gatan 4k x 4k K2 Summit direct electron detector. The Leginon software was used for data collection automation and resulting images were stored in the Appion database. Initial data processing was performed with cryoSPARC v3.2^64^, which included CTF correction using GCTF^65^, template picking, and 2D and 3D classification and refinement methods leading to a ∼3.3 Å C1 global reconstruction. The particles from this reconstruction were imported into Relion 3.1^66^, subjected to C3 symmetry expansion, followed by focused 3D classifications without alignments using a mask around the antibody Fab and S-protein RBD regions of a single protomer. Classes with well-resolved density in this region were selected and subjected to additional rounds of focused classification. Refinements were performed with limited angular searches and a mask around the trimeric S-protein and a single Fab. The final set of particles reconstructed to ∼3.7 Å global resolution.

Model building was initiated by rigid body docking of the x-ray structure of the Fab and a published cryo-EM model of the SARS-CoV-2 spike open state (PDB ID: 6VYB) into the cryo-EM map using UCSF Chimera^63^. Manual building, mutagenesis and refinement were performed in Coot 0.9.6^67^, followed by relaxed refinement using Rosetta Relax^68^. Model manipulation and validation was also done using Phenix 1.20^69^. More complete data collection, processing and model building statistics are summarized in Table S6. Figures were generated using UCSF ChimeraX^70^.

### Crystallization and X-ray structure determination

384 conditions of the JCSG Core Suite (Qiagen) were used for crystal screening of S728-1157 Fab crystals on the robotic CrystalMation system (Rigaku) at Scripps Research. Crystallization trials were set-up by the vapor diffusion method in sitting drops containing 0.1 μl of protein complex and 0.1 μl of reservoir solution. Crystals appeared on day 14, were harvested on day 21, pre-equilibrated in cryoprotectant containing 15% ethylene glycol, and then flash cooled and stored in liquid nitrogen until data collection. Diffraction quality crystals were obtained in solution containing 0.2 M di-Ammonium tartrate, 20% (w/v) polyethylene glycol (PEG) 3350. Diffraction data were collected at cryogenic temperature (100 K) on Scripps/Stanford beamline 12-1 at the Stanford Synchrotron Radiation Lightsource (SSRL). The X-ray data were processed with HKL2000^71^. The X-ray structures were solved by molecular replacement (MR) using PHASER^72^ with MR models for the Fabs from PDB ID: 7KN4^73^. Iterative model building and refinement were carried out in COOT^74^ and PHENIX^75^, respectively.

### Animals and challenge viruses

To determine whether mAbs in the panel could reduce viral load *in vivo*, Syrian hamsters (females, 6-8 weeks old) were intraperitoneally administered 5 mg/kg of candidate mAb 1 day after intranasal infection with 10^3^ PFU of SARS-CoV-2 viruses (an early SARS-CoV-2 isolate, Delta or BA.1 Omicron). Control animals were treated with an Ebola-specific mAb (KZ52) of matched isotype. At day 4 post-infection, lung tissues and nasal turbinate were collected to evaluate viral titers by standard plaque assay on Vero E6/TMPRRSS2 cells. The animal study was conducted in accordance with the recommendations for care and use of animals by the Institutional Animal Care and Use Committee at the University of Wisconsin under BSL-3 containment using approved protocols.

### Biolayer interferometry (BLI)

To determine precise binding affinity, the dissociation constant (K_D_) of each mAb was performed by biolayer interferometry (BLI) with an Octet K2 instrument (Forte Bio/Sartorious). The trimeric spike SARS-CoV-2 and its variants were biotinylated (EZ-Link Sulfo-NHS-Biotin, ThermoFisher), desalted (Zeba Spike Desalting, ThermoFisher), and loaded at a concentration of 500 nM onto streptavidin (SA) biosensor (Forte Bio/Sartorious) for 300 s, followed by kinetic buffer (1x PBS containing 0.02% Tween-20 and 0.1% bovine serum albumin) for 60 s. The biosensor was then moved to associate with mAbs of interest (142 nM) for 300 s, followed by disassociation with the kinetic buffer for 300 s. On rate, off-rate, and K_D_ were evaluated with a global fit, the average of those values with high R-squared from two independent experiments were presented. Analysis was performed by Octet Data Analysis HT software (Forte Bio/Sartorious) with 1:1 fitting model for Fabs and 1:2 interacting model for IgG.

For competitive assay by BLI, streptavidin (SA) biosensor was pre-equilibrated in 1xPBS for at least 600s to bind with the biotinylated trimeric spike WT-6P and spike BA.1 Omicron-6P for 300s. The first mAb was associated on the loaded sensor for 300s, followed by the second mAb for another 300s. The final volume for all the solutions was 200 μl/well. All of the assays were performed with kinetic buffer at 30°C. Data were analyzed by Octet Data Analysis HT software (Forte Bio/Sartorious) and plotted using GraphPad Prism.

### Statistics

All statistical analyses were performed using GraphPad Prism software (version 9.0). The numbers of biological repeats for experiments and specific tests for statistical significance used are described in the corresponding figure legends. P values of ≤ 0.05 were considered significant [*\*, P ≤ 0*.*05; **, P ≤ 0*.*01; ***, P ≤ 0*.*001; ****, P< 0*.*0001*), while P values of > 0.05 were considered as non-significant (ns)].

